# Optimized CRISPR guide RNA library cloning reduces skew and enables more compact genetic screens

**DOI:** 10.1101/2022.12.22.521524

**Authors:** Seok-Jin Heo, Lauren D. Enriquez, Scot Federman, Amy Y. Chang, Rachel Mace, Kaivalya Shevade, Phuong Nguyen, Adam J. Litterman, Shawn Shafer, Laralynne Przybyla, Eric D. Chow

## Abstract

The development of CRISPR genetic screening tools has improved functional genomics, as these tools enable precise genomic editing, provide broad access to genomic regions beyond protein-coding genes, and have fewer off-target effects than other functional genomics modalities, allowing for novel applications with smaller library sizes compared to prior technologies. Pooled functional genomics screens require high cellular coverage per perturbation to accurately quantify phenotypes and average out phenotype-independent variability across the population. While more compact libraries have decreased the number of cells needed for a given screen, the cell coverage required for large-scale CRISPR screens still poses technical hurdles to screen in more challenging systems, such as iPSC-derived and primary cells. A major factor that influences cell coverage is screening library uniformity, as larger variation in individual guide RNA abundance requires higher cell coverage to reliably measure low-abundance guides. In this work, we have systematically optimized guide RNA cloning procedures to decrease bias. We implement these protocols to demonstrate that CRISPRi screens using 10-fold fewer cells than the current standard provides equivalent statistically significant hit-calling results to screens run at higher coverage, opening the possibility of conducting genome-wide and other large-scale CRISPR screens in technically challenging models.

## INTRODUCTION

The human genome project produced the first assembled human genome over 20 years ago (1, 2). Genomic sequencing efforts reveal genes and genetic variation associated with disease but for the most part do not reveal gene function. As such, functional genomics efforts have been critical to assign function to the roughly 20,000 human protein-coding genes identified. In the past decade CRISPR (Clustered Regularly Interspaced Short Palindromic Repeats)-based screens have increased the ease of genome-wide genetic screens, allowing researchers to find new components of biological pathways, assign mechanism to existing drugs, identify novel therapeutic targets, and uncover synergistic genetic relationships (3–7). However, due to the size of genome-wide guide libraries (20,000-200,000+ elements) and typical cell coverage (500-1000x) required, each screen typically requires tens to hundreds of millions of cells per sample (8–12). This requirement poses a logistical challenge to screening cell models where large-scale culturing is difficult, such as adherent cell lines, or growth-limited models such as primary and differentiated cell lines (13–15).

In this work, we report optimizations to several steps in CRISPR guide library cloning that significantly decrease guide representation bias, allowing for screening at lower cell coverage (16). Improvements were made in the following areas. First, ordering guide oligos in both forward and reverse complement orientations to counteract sequence-specific biases in oligo synthesis (17). Second, decreasing the number of PCR cycles used to prepare inserts to avoid over amplification to maintain library uniformity. Lastly, working at low temperatures during insert preparation to reduce biased dropout of inserts with lower melting temperatures (T_m_). We used these optimizations to clone new versions of published genome-wide CRISPRi and CRISPRa libraries(18) and achieved more uniform guide distributions, as evidenced by reduced skew ratios compared to the legacy libraries available on Addgene (#83969 and #83978; referred to as legacy throughout the text).

With the improved CRISPRi library, we demonstrate comparable performance in survival screens at 100 versus 1000-fold coverage. In a survival screen coupled with treatment with the tyrosine kinase inhibitor dasatinib in K562 cells, we observe more hits in expected pathways compared to those identified in parallel screens run using a publicly available CRISPRi library under more stringent analysis conditions. Lastly, through a transduction titration experiment, we demonstrate the feasibility of performing screens at 50-fold cell coverage, facilitating genome-wide screens requiring only 5 million cells per sample for a 100,000-guide library. This level of coverage will enable researchers to use more sophisticated and biologically relevant readouts such as live cell FACS-based approaches and model systems such as adherent cells, iPSC-derived cells, and primary cells that were previously challenging or impossible to work with at genome-scale. Moreover, the cloning methods described here are generalizable across guide libraries generated for any type of CRISPR based approach including nuclease, base editing (19), prime editing (20), CRISPRoff (21), and RNA editing applications, and are applicable across a range of Cas enzymes including Cas9, Cas12, Cas13, and others.

## MATERICALS AND METHODS

### Cell Lines

ECCAC K562s were cultured according to standard protocols and transfected with lentivirus containing the dCas9-KRAB construct. Cells were then sorted using the BD FACS Aria based off BFP signal. The pooled cell line was utilized for the essential gene drop-out screens, transduction experiments, and dasatinib screens. Cell lines were maintained in shaking cultures at 100 rpm at a concentration of 500,000 cells / mL for experiments.

### Plasmid Vectors

The lentiviral expression vectors used in pooled CRISPR screen experimentations are available on Addgene under the following names: pLGR1002, Addgene 188320.

### Reagents

Dasatinib was acquired from Millipore Sigma (Catalog Number SML2589). The dasatinib was diluted with DMSO to final concentration of 4 μM. Drug dosage for K562s was determine by performing a drug titer test and a cell viability curve was generated. A single dose was applied to K562s and cells remained in treated media for 72 hours.

### Guide library cloning

Detailed protocol is in the supplemental materials. The main highlights are ordering insert oligo pools in both orientations, minimizing over amplification of the insert, and performing gel electrophoresis size selection and extraction at low temperatures.

### Plasmid Library Virus production and titers

Guide library virus was prepared using protocols from the Weissman Lab (https://weissman.wi.mit.edu/crispr/) with Lenti-X 293Ts (Catalog Number 632180). Virus titers were performed with polybrene (Catalog Number: 3597560) in the K562 cell lines and Multiplicity of Infection (MOI) and percent infection were determined by BFP signal using flow cytometry.

### Essential Gene drop-out screens

K562 cells were infected at ∼25%, targeting a 100 or 1000-fold cell coverage. After two days, transductants were selected with puromycin for 3 days until approximately 90% of the culture was BFP positive. The cultures were maintained at 500,000 cells / mL density for 7 days. Cell pellets were collected and frozen before DNA extraction. A schematic of the screen timeline is illustrated in Supplementary Figure S6A.

### Dasatinib screens

Approximately 10 million K562s with the dCas9-KRAB constructs were infected with either the LGR or legacy CRISPRi V2 libraries containing the top 5 guides (resulting in approximately 100-fold cell screening coverage). Puromycin treatment lasted for 5 days (until approximately 90% enrichment). 7 days after infection the T_0_ samples were collected for each library. A single dose of 0.75 nM of dasatinib was use as the drug selective pressure and 0.01% DMSO was used as the control. Treatment was for 72 hours with 6 days of recovery. A schematic of the screen timeline is illustrated in Supplementary Figure S6C. The LGR and legacy screens experienced similar cell death and recovery growth (Supplementary Figure S9A).

### Genomic DNA Processing

Genomic DNA was extracted using Macherey-Nagel Nucleospin Blood kits. The 10x cell pellets were processed with the Mini kit (Catalog Number: 740951), the 50x and 100x pellets with the L kit (Catalog Number: 740954), and the 200x and 1000x pellets with the XL kit (Catalog Number: 740950). All pellets were processed according to the kit-specific protocols and quantified by Nanodrop.

### NGS sample prep and sequencing

Oligo pool libraries were performed with the Claret Bioscience SRSLY PicoPlus kit (K250B-24) according to manufacturer instructions with 20-25 ng oligo template and 10 PCR cycles. Libraries were pooled and sequenced on a NextSeq 550.

The gDNA samples were prepared for NGS sequencing through a process of PCR amplification, PCR sample pooling, double-sided SPRI bead purification, quantification, and NGS library pooling. 24 cycles of PCR were performed using NEBNext Ultra II Q5 Master Mix (M0544). The forward primers (5’ PCR primers) used in the NGS library preparation for gDNA samples had the Illumina 5’ adapter with a 6 bp index barcode (for de-multiplexing pooled samples): AATGATACGGCGACCACCGAGATCTACACGATCGGAAGAGCACACGTCTGAACTCCAGTCACnnnnnnGCACAAAAGGAAACTCACCCT. The common reverse primer (common 3’ PCR primer with the Illumina 3’ adapter sequence) used in the NGS library preparation for sgRNA libraries cloned into the pLGR1002 lentiviral expression vector:

CAAGCAGAAGACGGCATACGAGATATGCTGTTTCCAGCTTAGCTCTT. The common reverse primer used for the legacy libraries is: CAAGCAGAAGACGGCATACGAGATCGACTCGGTGCCACTTTTTC. Due to different T_m_ of the reverse primers, the LGR PCR samples were amplified with an annealing temperature of 62.6°C, while the legacy samples were amplified with an annealing temperature of 65°C. These cycle number and temperature conditions were optimized based off conditions previously established by the Weissman Lab. All PCRs were conducted in 100 μL volumes with 10 μg of DNA per reaction. After PCR, aliquots of each set of reactions were pooled. The legacy samples were purified using 0.65x and then 1x doubled-sided SPRI beads while the LGR samples using 0.65x and then 1.2x double-sided SPRI beads. The purified and pooled LGR and legacy samples were quantified on the TapeStation using the High Sensitivity Kit before being sequenced on the NextSeq 550 with a custom sequencing primer (GTGTGTTTTGAGACTATAAGTATCCCTTGGAGAACCACCTTGTTGG). PhiX control library was spiked in at 10% PhiX to increase base diversity for the single-end 20 cycles sequencing runs.

Note that the first several cycles of sequencing on the NextSeq 550 are used to identify clusters. Because G-bases are dark in two-color sequencing, sequences that contain a polyG sequence are difficult or impossible to identify. This issue will also be present on the MiniSeq platform. Two-colored patterned flow cell systems such as the NovaSeq and NextSeq 1000/2000 should not be affected by this. An alternative method to increase compatibility with the NextSeq 500/550 is to use a staggered sequencing approach.

### Data Availability/Novel Programs, Software, Algorithms

The volcano plots were generated using ScreenProcessing (https://github.com/ucsf-lgr/ScreenProcessing). ScreenProcessing (5, 22) analyzes pooled CRISPR screens by comparing the sgRNAs targeting each gene of interest with the entire set of sgRNAs targeting all genes. sgRNAs are ranked according to their enrichment score, which is the comparison of phenotype distributions of sgRNAs targeting each gene of interest with the negative control sgRNAs (NTCs) that target non-coding genes. The NTCs serve as a reliable Null Distribution. Genes are ranked according to the phenotype scores and p-values derived from the sgRNA rankings. The phenotype score is the average of the top 3 sgRNAs targeting the gene and is calculated using the absolute value of log_2_ enrichment score (22). The p-value is calculated using the Mann-Whitney U test (MW test) by comparing the top 5 sgRNAs to the NTCs. A gene is considered a hit (with a False Discovery Rate of <0.05) if the | effect size Z-score (Absolute Phenotype) x log_10_p-value| ≥ 20. ScreenProcessing visualizes this data as a volcano plot, plotting the phenotype score verses the log_10_p-value, and labels genes, gene hits, NTCs, and NTC hits accordingly.

Quality Control plots as well as gene hit analysis were done with MAGeCK-Vispr (23, 24). MAGeCK-Vispr uses the Negative binomial p-value to perform sgRNA ranking and the Expectation Maximization (EM) Algorithm to perform gene ranking. The false-discovery rate (FDR) can be adjusted on the experiment web interface that MAGeCK generates. Gene Onology (GO) and KEGG pathway analysis for the dasatinib screens were performed with clusterProfiler (25–28) and DAVID (29, 30).

## RESULTS

### Cloning optimization reduces library bias

To improve guide cloning efficiency, we performed a series of optimizations to improve the representation of CRISPR sgRNA libraries using sequences from previously described genome-wide CRISPRi and CRISPRa libraries (18). To clone these libraries, single stranded oligo templates are amplified and converted to double stranded inserts that are cloned into a vector. In the CRISPRi/a library cloning protocol, the insert is digested into a 33bp double-stranded product, gel purified and ligated into a lentiviral expression vector (pLGR1002). We first examined whether the polymerase used to synthesize double stranded DNA encoding the sgRNA insert could impact library representation. In a pilot experiment comparing three different polymerases (Klenow, Klenow exo-, and NEB Q5 Ultra II), we observed varying guide representations in a library of 192 sgRNAs, (Figure 1A) with the most uniform representation observed in inserts prepared with Q5 Ultra II polymerase. To check if the clones contained the expected insert, we performed colony PCR using PCR primers outside of the BstXI and BlpI restriction sites (Supplementary Figure S1A). We observed that the majority of individual clones in libraries prepared with all three polymerases had the expected 290bp band but 40% also exhibited higher molecular weight bands (Supplementary Figure S1B). Sanger sequencing of plasmids minipreps from these colonies showed mixed bases in the spacer region of the sgRNA (Supplementary Figure S1C). We hypothesize that these mixed sequences arise from transformation with plasmids containing non-complementary hybrid inserts. As predicted from this model, retransformation of these plasmids yielded colonies that gave a single band upon colony PCR (Supplementary Figure S1D).

**Figure 1.**
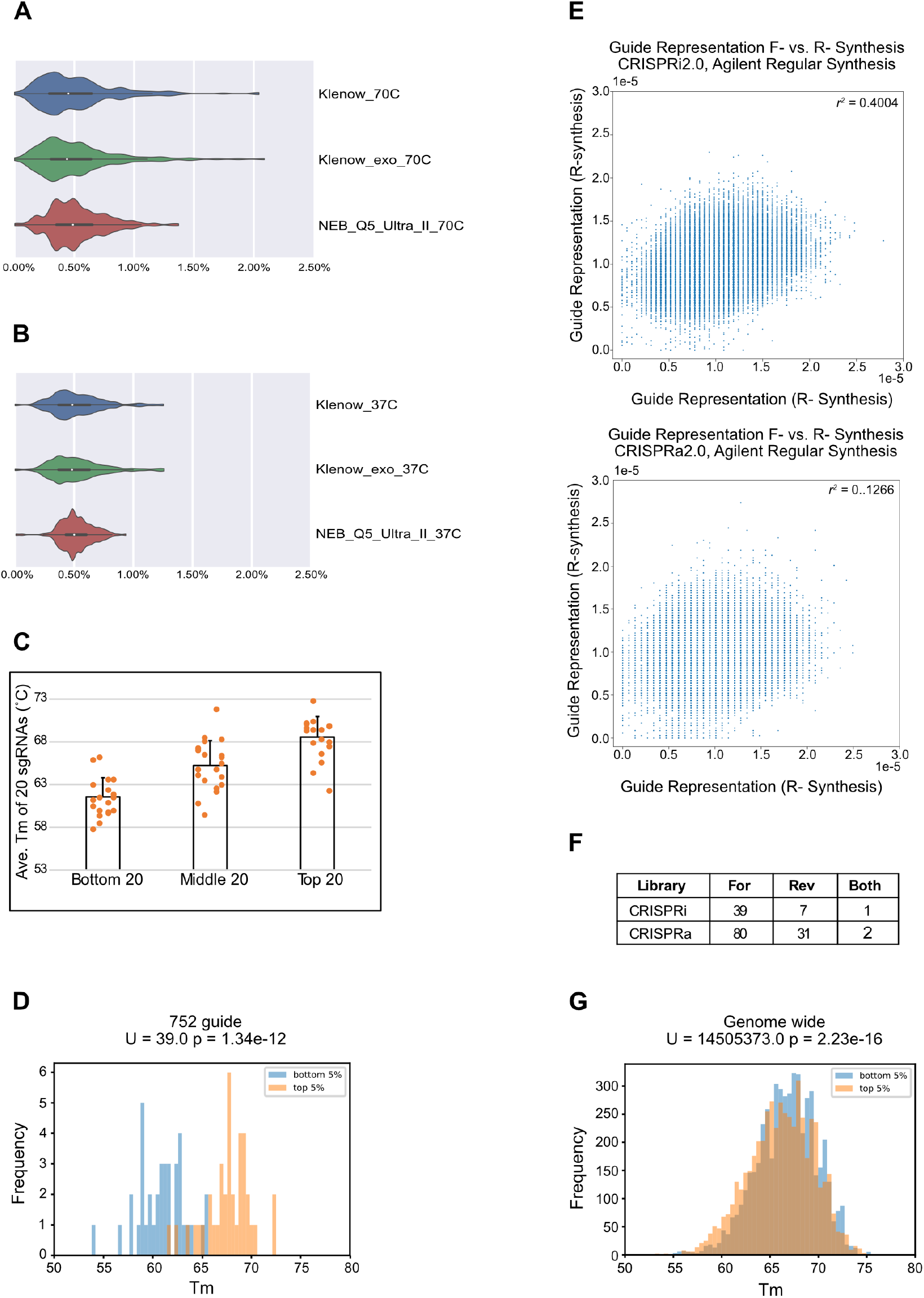
Factors affecting guide cloning uniformity. (A) Violin plots depicting guide abundance distributions of libraries prepared with three different polymerases (Klenow, Klenow exo-, and NEB Q5 Ultra II) and inserts extracted at 70°C. (B) Violin plots of libraries prepared the three different polymerases and a 37°C extraction. (C) Melting temperature (Tm) and abundance of the lowest, mid, and top 20 guides in the 752 element pilot library. (D) Mann-Whitney U-test comparing T_m_’s of high (top 5%) and low (bottom 5%) abundance guides in the 752 element pilot library. (E) Correlation between forward and reverse complement oligo pools for CRISPRi V2 (top) and CRISPRa V2 (bottom). (F) Number of guides missing from the oligos pools in forward, reverse, or combined sets in the CRISPRi V2 and CRISPRa V2 guide libraries. (G) Mann-Whitney U-test comparing the T_m_’s of high (top 5%) and low (bottom 5%) abundance guides in the CRISPRi V2 genome-wide library.

Next, we tested if a lower 37°C insert elution during gel purification narrows guide distribution and reduced the formation of hybrid clones. Using the same starting material, we found that the lower elution temperatures increased uniformity and Q5 Ultra II polymerase still performed better than Klenow (Figure 1B). Furthermore, the lower elution temperature reduced the formation of hybrid clones from 40% to 1.2% (Supplementary Figure S2). We also observed that PCR amplification of oligo pools produced a single narrow band of product, whereas primer extension with Klenow yielded non-specific products that were both smaller and larger than the intended insert (Supplementary Figure S3).

We moved on to a larger pilot library of 752 guide RNAs to further optimize the cloning protocol. Since Q5 Ultra II polymerase performed better than the two different mesophilic polymerases, we sought to determine the effect of additional PCR cycles on guide distribution. We repeated the library cloning described above with either one or fifteen cycles of insert PCR. A pairwise comparison indicated similar representations of the 752 gRNAs across the two libraries (Supplementary Figure S4). During this experiment, we observed higher molecular weight smears in some PCR products (Supplementary Figure S5A) that are likely overamplification products that could contribute to the hybrid clones observed earlier. We reduced PCR cycle number and optimized template concentration to minimize production of nonspecific products (Supplementary Figure S5B). In this new library, even though the gel-purified insert was eluted at 37°C, we still observed a relationship between guide abundance and the melting temperature (T_m_) of the inserts (Figure 1C). The average T_m_ of the 20 most highly represented gRNAs was higher than 68°C. In contrast, the average melting temperature of the 20 most lowly represented gRNAs was less than 62°C. A Mann-Whitney U test comparing the lowest and highest 5^th^ percentiles of guide representation (n_1_ = 38, n_2_= 38) indicated a statistically significant difference between the T_m_ distributions of these two populations (Figure 1D). These results indicate that a 37°C elution temperature can still result in guide bias due to T_m_.

We next sought to reduce heterogeneity of guide abundance in the template oligo pool. Since oligo sequence can affect yields, we ordered oligos encoding the genome-wide CRISPRi and CRISPRa libraries in both the forward and reverse complement directions (F, R), hypothesizing that this would minimize effects of sequence-specific oligo synthesis bias. To test this, we prepared NGS libraries from the oligo pools with a single stranded DNA library preparation kit. In comparing the representation of guides derived from either the F- or R-strand, we observed a weak correlation between the representation of guide sequences derived from different strand synthesis pools (Figure 1E) for both the CRISPRi and CRISPRa libraries. Additionally, we observed a non-overlapping subset of guides missing from oligos synthesized in either orientation (Figure 1F). These results indicate that ordering oligos in both the forward and reverse complement orientations increase guide uniformity and minimizes guide dropout.

To test the improved cloning strategy, we cloned two genome-wide guide libraries using our improved protocol. Since a T_m_-dependent guide bias was still present with a 37°C elution, we performed insert gel electrophoresis on ice and reduced the elution temperature to 4°C. We recloned the V2 CRISPRi and CRISPRa libraries which respectively contain 103,073 and 101,242 elements, (18). After sequencing the cloned libraries, we repeated the Mann-Whitney U test comparing the melting temperature of the lowest 5% represented guides (n = 5151) against the highest 5% represented guides (n = 5151) and observed a ρ statistic of 0.547 (Figure 1G), compared to a ρ statistic of 0.025 for 752-guide library prepared with a 37°C elution. The ρ statistic values indicate that the lowest 5% most represented guides and the highest 5% represented guides shared a similar underlying distribution, which was not the case in the 752-guide dataset used insert templates synthesized in a single orientation and a 37°C insert elution temperature.

When compared to the legacy libraries, both of our libraries show a more uniform distribution (Figure 2A, 2B) with fewer dropouts. Skew ratios used to represent library uniformity are calculated by comparing the abundance of guide pairs at different percentiles, with lower ratios indicating more uniform libraries. In a 100,000 element library, a 90/10 skew ratio is calculated by comparing the abundance of the 10,000^th^ top and bottom elements. Our libraries have a 90/10 skew ratio under 2, outperforming the legacy libraries as well as a commercially available CRISPR knockout library (https://go.takarabio.com/genomewidelibrary.html). The uniformity is more evident when comparing skew ratios at the extremes of the distribution (Figure 2C).

**Figure 2.**
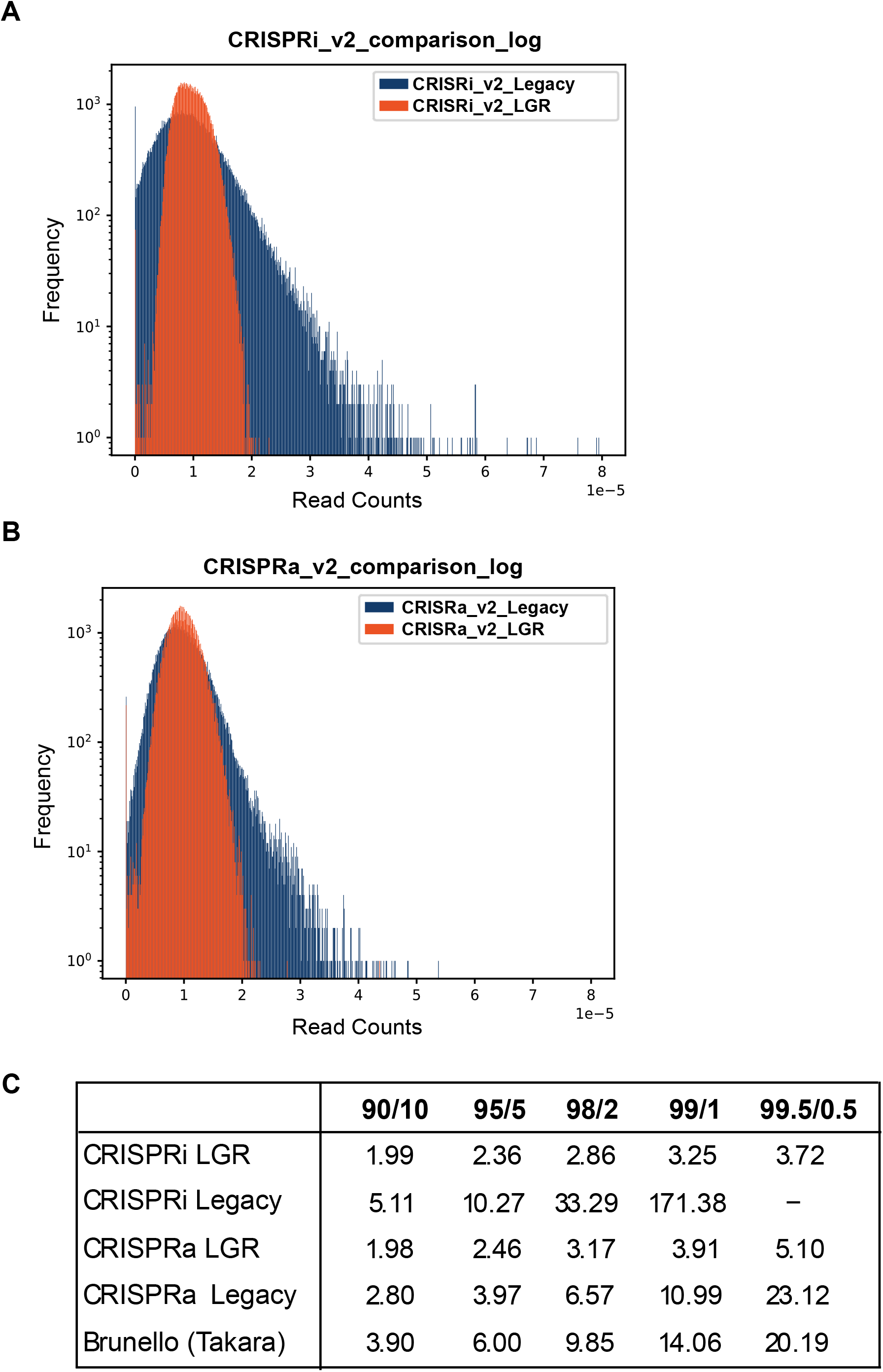
New optimizations in the cloning protocol improve genome-wide guide libraries. (A) Histograms of the CRISPRi V2 legacy library (blue) compared to the optimized CRISPRi V2 LGR library (orange) show that the LGR library has a tighter distribution of sgRNAs. (B) Similarly, the CRISPRa V2 LGR library (orange) shows a tighter distribution than the CRISPRa V2 legacy library (blue). (C) Skew ratio table comparing CRISPRi V2 and CRISPRa V2 LGR libraries with several publicly available libraries.

### Lower skew library performs well at lower cell coverage

Previous work suggested libraries with 90/10 skew ratios below 2 could be screened at 100x cell coverage (31). To test this, we performed a genome-wide CRISPRi survival screen in K562 cells expressing dCas9-KRAB transduced and maintained at 100x or 1000x guide library coverage (Figure 3A; Supplementary Figure S6A). Cells were transduced ranging from 20.2 – 23.7% infection to minimize cells containing multiple integrations. We compared the essential genes identified in our 1000x screen to those previously identified (18). The original study used the expanded library of 10 guides per gene. To perform the comparison, we extracted data from the top 5 guides for each gene and analyzed the data with the ScreenProcessing pipeline. Essential genes are identified by the gamma score, which represents the growth enrichment (log_2_ enrichment) determined by sgRNA read counts between the untreated sample and T_0_. In Figure 3B, the volcano plots for each screen shows the essential genes on the left side of the volcano plots (labeled as gene hits). The Horlbeck et al. 2016 screen identified 1,883 essential genes, whereas our new library screen identified 1,802 essential genes (Figure 3B). Between the LGR 1000x and Horlbeck et al. 2016 1000x screens, there is an overlap of 1,369 essential genes (Figure 3C). Some discrepancies in essential genes identified by each screen were expected due to differences in user handling, reagents, and cell doublings (∼8 doublings in our screen compared to ∼10).

**Figure 3.**
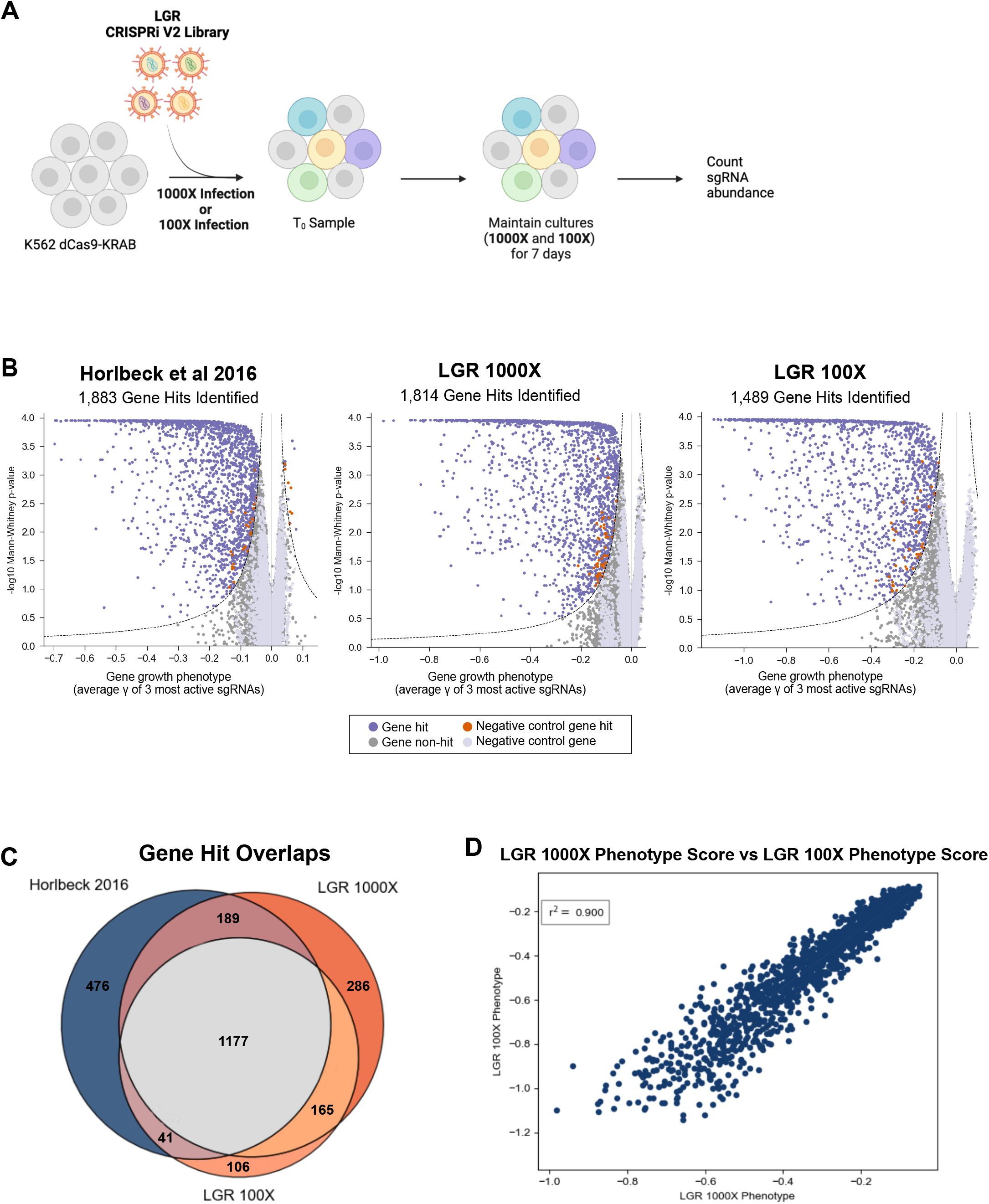
The optimized LGR library performs similarly to the existing legacy library, even at lower cell coverage. (A) Schematic of the CRISPRi V2 survival screen in K562 cells performed with the LGR library at 100 or 1000-fold cell coverage. (B) Comparison of the 1000-fold cell coverage screen performed by Horlbeck 2016 using the CRISPRi V2 legacy library (left) versus the 1000-fold (middle) and 100-fold (right) cell coverage screens performed using the CRISPRi V2 LGR library. (C) A diagram illustrating the amount of overlap in essential genes identified in each screen. (D) A point comparison of the phenotype score of overlapping hits between the CRISPRi V2 LGR 100 and 1000-fold screens.

There was greater overlap between the 1000x and 100x screens using our new library (Figure 3C). This is expected since the same library and cell doublings were used. The 100x screen identified 1,489 essential genes (Figure 3B). To compare the quality of the 100x and 1000x screen, we compared the phenotype scores for common gene hits. The pairwise comparison plot of the phenotype scores for gene hits shows a coefficient of determination of 0.900 (r^2^) using a linear least-squares regression (Figure 3D). This demonstrates that our new library generates very similar gene-level results between 1000x and 100x coverage survival screens. While there was not perfect overlap between all three screens, the unique hits in each screen tended to fall near the cutoff (Supplementary Figure S7B), populated by weaker and/or less significant hits.

The strong correlation between the 100x and 1000x screens suggested we might be able to screen libraries at even lower coverages. To test this, we performed transductions at 200x, 100x, 50x, and 10x coverage in technical duplicate using both the LGR and legacy libraries. The percent infection for these samples ranged from 11.5 – 22.8% and cells were treated with puromycin for 5 days until transduced cells accounted for approximately 90% of cells (Supplementary Figure S6B). T_0_ samples were collected at this point and processed for NGS (Figure 4A). Both libraries maintained high correlations of guide representation above 0.8 at all coverages 50x or greater (Figure 4B). However, the LGR library maintained lower skew ratios at all tested coverages (Figure 4C). Furthermore, the 50x and 100x samples showed similar skew ratios to the 200x sample. In contrast, the legacy library skew ratio was worse at 100x and even worse at 50x (Figure 4C). One concern with running screens at lower cell coverage is guide dropouts. Our new library showed similar rates of guide dropouts between 200x down to 50x while the legacy library showed increased dropouts from 200x to 100x and 50x. Notably, the majority (∼94.62%) of sgRNA sequences dropped from the LGR library began with a PolyG sequence (Supplementary Figure S8A) and the dropout was likely due to a technical artifact of sequencing on the 2-color NextSeq 550 (See technical note in protocol). This means the true number of dropouts in the 50x samples could be as low as 6-7 guides. In contrast, sgRNA sequences that began with the PolyG sequence only accounted for a minority of the dropouts in the legacy library (Supplementary Figure 8A). Furthermore, re-sequencing the plasmid libraries on the HiSeq 4000, a system not susceptible to polyG sequences at the beginning of the read resulted in dropout of only two guides and a lower skew ratio for our new CRISPRi library (Supplementary Figure S8B). The low skew ratio and sgRNA dropout number for our library suggests this library can be used in screens with as little as 50x cell coverage, ∼5 million cells for a 100,000 element genome-wide library.

**Figure 4.**
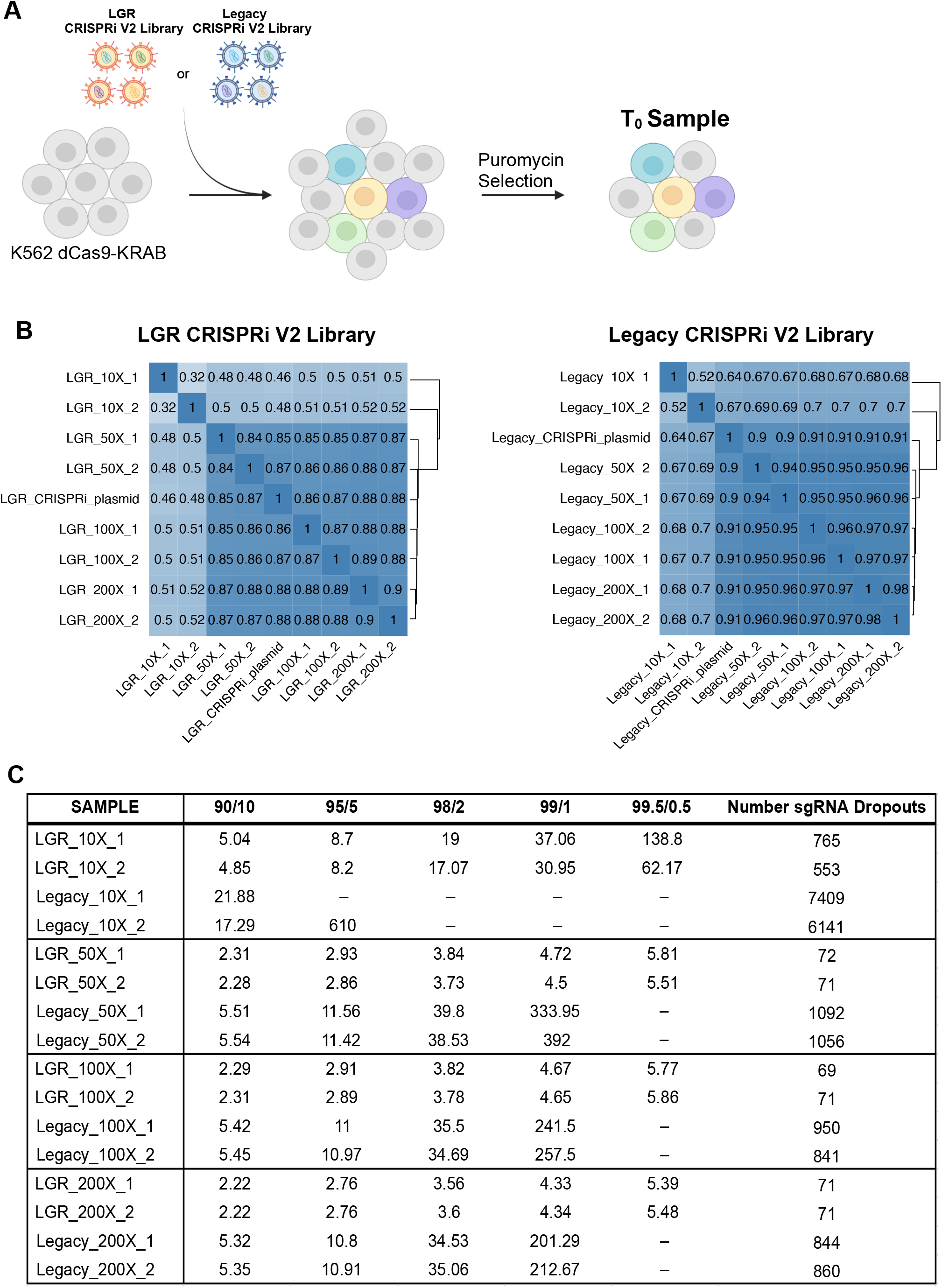
Transduction titration comparisons between the CRISPRi V2 LGR and the legacy libraries. (A) Schematic of the transduction experiments performed using the LGR and legacy libraries at 10, 50, 100, and 200-fold cell coverage. Each library at each coverage had a biological replicate. (B) Hierarchical clustering of transduced guides with the LGR library (left) and the legacy library (right). (C) Guide library skew ratios and number of sgRNA dropouts in the LGR and legacy samples at 10, 50, 100, and 200-fold cell coverage.

Although these screening experiments demonstrated promising results, they were simple survival screens. To understand the performance of our library in a screen with selective pressure, we performed a drug survival screen on K562, a chronic myeloid leukemia (CML) cell line using dasatinib at 100x coverage to determine whether a high-quality screen can be performed at this low coverage (Figure 5A). Dasatinib is a broad-spectrum tyrosine kinase inhibitor (TKI) that is approved for the treatment of CML (32–36). However, there is no cure for CML as 40% of cases of clinical TKI failure occur in the setting of sustained BCR-ABL1 inhibition (37). Identification of genes and biological pathways that are synthetically lethal to CML or CML plus dasatinib could result in new CML therapies (38). Given the performance of the LGR library at lower coverages, we hypothesized that a 100x coverage screen could be performed to comprehensively identify genes related to dasatinib resistance and sensitivity and demonstrate that lower coverage screens in other challenging systems could be performed with our improved library.

**Figure 5.**
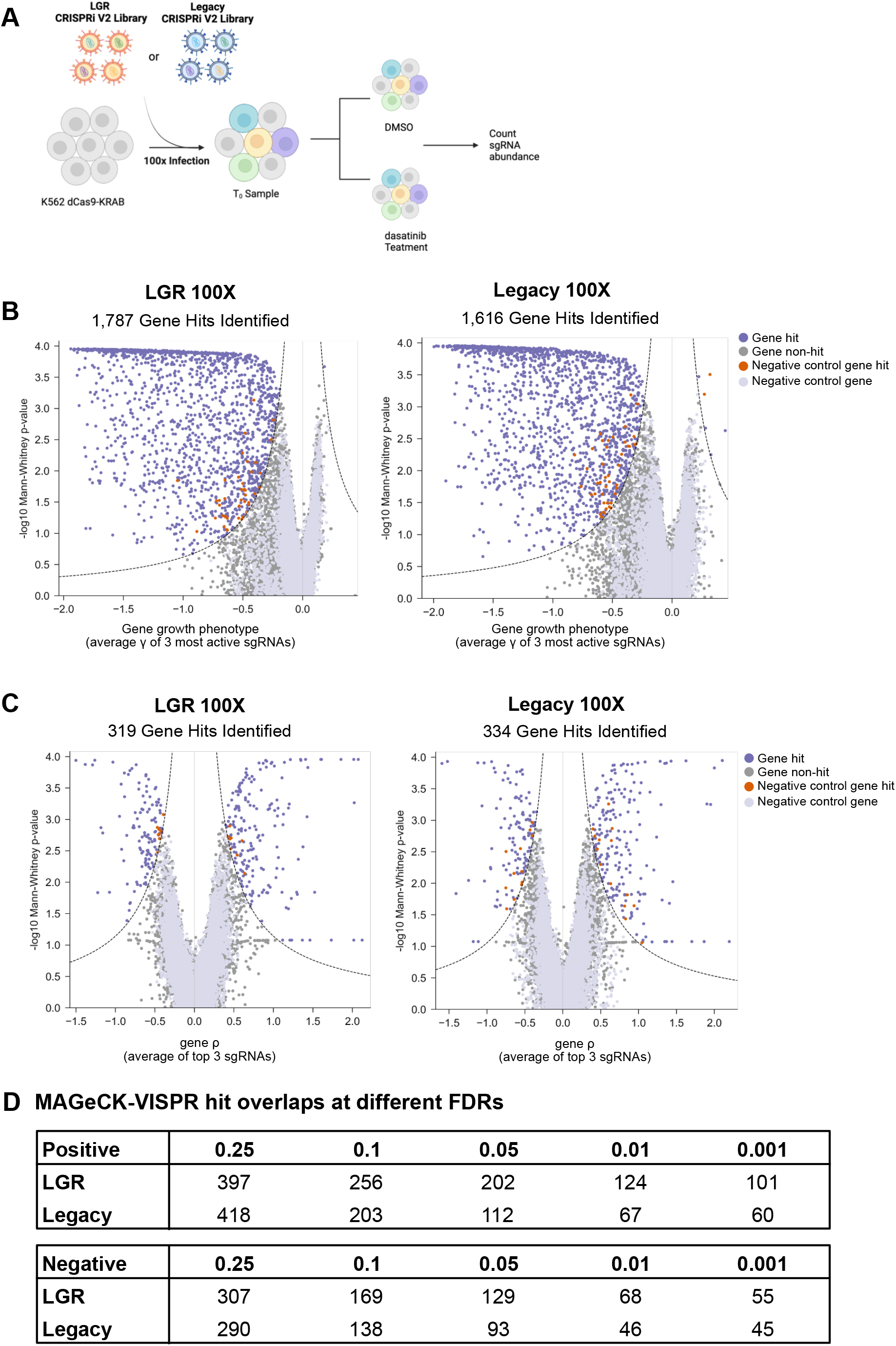
The CRISPRi V2 LGR library identifies more bonafide hits in a 100-fold cell coverage K562 dasatinib survival screen. (A) Schematic of the dasatinib survival screen performed using the LGR and legacy libraries at 100-fold cell coverage. (B) Comparison of essential gene hits (T_0_ versus DMSO samples) identified in the LGR (right) versus the legacy(left) libraries. (C) Comparison of dasatinib treatment hits (dasatinib treatment versus DMSO control samples) identified in the LGR (right) versus the legacy (left) libraries. (D) MAGeCK-VISPR was used to determine the number of gene hits identified in legacy (left) libraries. (D) MAGeCK-VISPR was used to determine the number of gene hits identified in (increased survival) or negative (decreased survival).

### Genome-wide CRISPR dug screen yields more hits with new library

The dasatinib survival screens were performed at 100-fold cell coverage for both CRISPRi libraries in parallel (Figure 5A and Supplementary Figure S6C). Each library had one transfection in K562s and samples collected from each library starting from T_0_ were treated as technical replicates. The LGR and legacy libraries had 20% and 18% infection levels respectively. The samples were treated with puromycin for 5 days (until approximately 90% enrichment) and 7 days after infection the T_0_ samples were collected for each library. Each library had four technical replicates, two treated with 0.75 nM of dasatinib (single dose) and two treated with 0.01% DMSO (vehicle control) for 72 hours (Supplementary Figure S9A). After 72 hours, the samples were resuspended in fresh media and allowed 6 days to recover.

Dasatinib selective pressure reduced cell viability as expected and after removal of drug, cultures recovered (Supplementary Figure S9A). The LGR and legacy library samples (T_0_, DMSO, and dasatinib) clustered by library in the quality control heat map (Supplementary Figure S9B). The PCA plots showed a separation of the libraries in PC1 whereas PC2 and PC3 were driven by biological conditions (Supplementary Figure S9C). PC2 illustrates the effect of cell growth for 8 days after T_0_ and PC3 reflects the dasatinib treatment (Supplementary Figure S9C). From the DMSO vehicle control arm, more essential genes were identified with the LGR library (1,787) than the legacy library (1,616) (Figure 5B). Dasatinib-specific gene hits were identified by ScreenProcessing as rho, which represents the growth enrichment (log_2_enrichment) determined by sgRNA read counts between the treated sample (dasatinib) and the untreated sample (DMSO) (Figure 5C). Similar number of genes were identified by both libraries. Next, we used MAGeCK-Vispr (23, 24) to identify screen hits at various false-discovery rates (FDR) (Figure 5D). At the highest FDR of 0.25 the legacy and LGR libraries had similar numbers of total gene hits, 708 versus 704, respectively. However, as stringency increased, the LGR library yielded more gene hits for both positive (increased cell survival) and negative (decreased cell survival) categories. At the most stringent FDR used (0.001), the LGR CRISPRi V2 library had a total of 156 gene hits whereas the legacy library had 105.

The dasatinib gene hits for each library at FDRs of 0.25 and 0.001 were analyzed across the Gene Ontology (GO) and Kyoto Encyclopedia of Genes and Genomes (KEGG) databases to evaluate the quality of the hits generated (39–42). The mediator complex (GO ID: 0016592) and the oxidative phosphorylation pathway (KEGG ID: hsa00190) were identified among the most significantly enriched annotations for both libraries (Supplementary Figure S10). The mediator complex and the oxidative phosphorylation pathway have been shown to be potential drug targets to synergize with tyrosine kinase inhibitor treatments in CML. TKIs primarily target differentiated cells and fail to eliminate leukemic stem cells (LSCs) (43, 44). However, inhibiting mitochondrial oxidative phosphorylation in combination with TKI treatment eliminates LSCs (45). On the contrary in a genome scale CRISPR knockout screen it was shown that knocking out components of the mediator complex provided resistance against TKI treatment (46). Although the prior study used a CRISPR knockout library, several studies have shown strong correlation between CRISPR knockout and CRISPRi screens (47, 48)

In agreement with these two studies, we observe depletion of guides targeting the components of oxidative phosphorylation pathway and enrichment of guides targeting the mediator complex in the dasatinib screen. Our new library generated more hits for the mediator complex and the oxidative phosphorylation pathway at an FDR of 0.25 and 0.001 (Figure 6A). Additionally, the p-values of enrichment in both gene categories at both FDRs were markedly lower for the LGR library. Of the 40 genes annotated to be part of the mediator complex in the GO database the screen with our library identified 17 of those genes (including all 7 of the legacy library hits) at an FDR of 0.001 (Figure 6A, 6B). Additionally, our 100x screen identified more components of the mediator complex (17 vs 10) than the previous CRISPR knockout screen performed at 250x coverage (46). Furthermore, the 17 hits in our screen were identified at a lower FDR of 0.001. Only one hit passed that threshold in the prior study. The combination of our survival screen, transduction titration experiments, and drug perturbation screen demonstrate the feasibility of screening at much lower cell coverages with our improved library.

**Figure 6.**
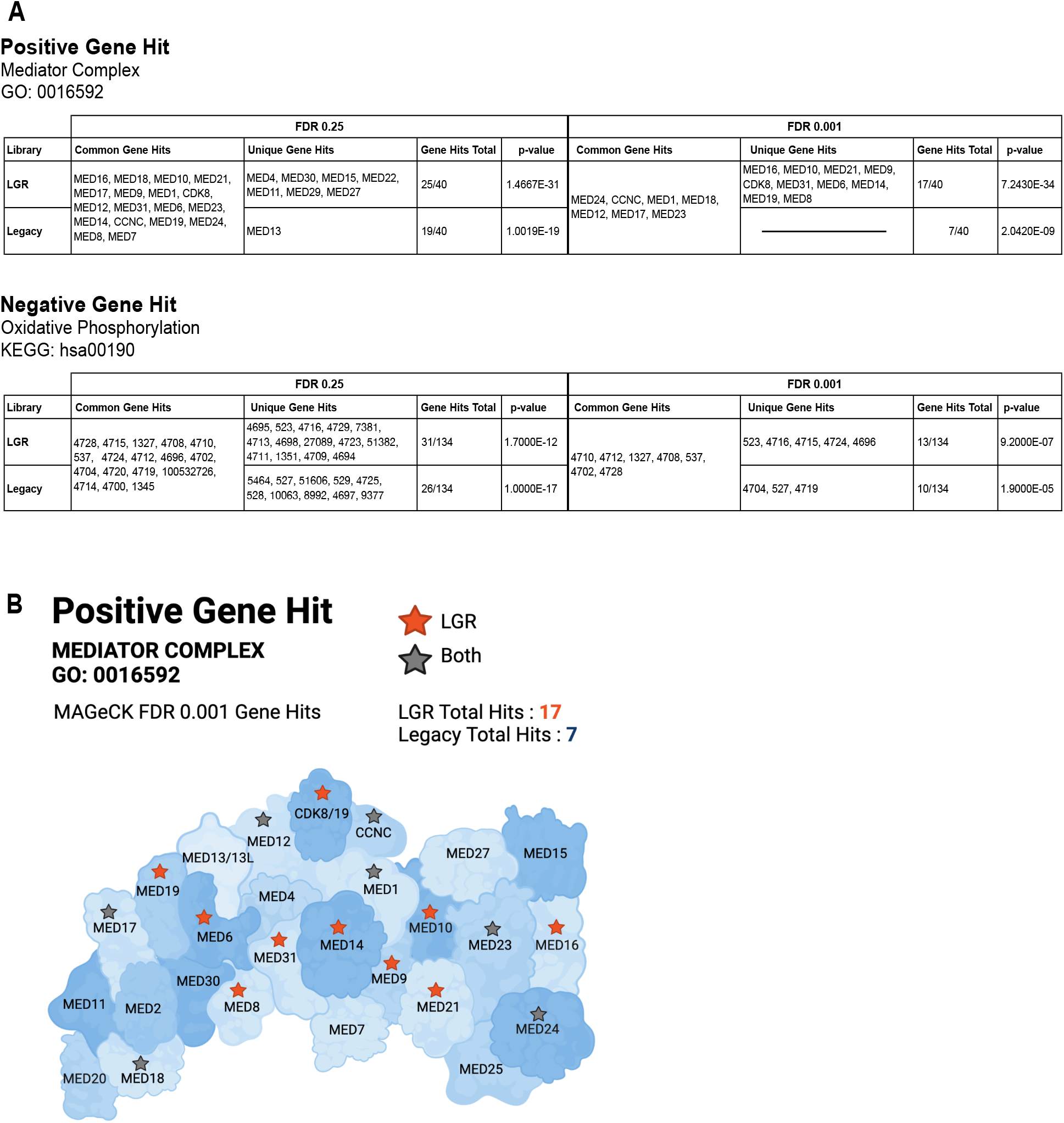
The top positive and negative gene hits in the dasatinib survival screens were investigated at 0.25 and 0.001 FDRs. (A) The list of gene hits determined by the LGR and legacy libraries for the Mediator Complex (top) and Oxidative Phosphorylation pathway (bottom). The Mediator Complex has a total of 40 genes associated with the cellular component and the Oxidative Phosphorylation pathway has 134. (B) Visual representation of hits in the Mediator Complex at a 0.001 FDR. LGR unique gene hits are marked by orange stars (total 10) and common gene hits (shared by the LGR and legacy libraries) are marked as grey stars (total 7).

## DISCUSSION

In this work, we have developed an improved guide library cloning method that can be applied to other libraries that are generated using oligo-derived sequences. Through a combination of ordering oligo templates in both forward and reverse complement orientation, optimizing insert amplification, and minimizing temperature during insert preparation steps, we have generated very uniform libraries that allow lower cell coverage screens. This has several practical benefits.

First, starting with the same number of cells and same library size, the new protocol enables screens with 10-20 times more samples. These could be biological replicates, additional perturbations, additional cell lines or clones, or isogenic controls (49). For example, more essential genes were identified in the 100x genome-wide CRISPRi survival screen in K562 when more replicates were analyzed (Supplementary Figure S11). This additional data can improve the quality and scope of screens. Second, if the same number of cells are used, a 2 million element library could be screened instead of a library containing 100,000 elements. This enables screens with larger libraries such as tiling screens to identify regulatory regions in non-coding sequences or synthetic combinatorial libraries. Third, due to the large number of cells that must be maintained in higher coverage screens, researchers often must split cells every day for several weeks. With lower cell coverage, cultures can be diluted further while still maintaining adequate coverage and split every two or three days if large tissue culture vessels are used (Supplementary Figure S12). Lastly, the majority of CRISPR screens have been performed in transformed cell lines because their cultures can easily be scaled up. This has been adequate for certain areas of biology such as cancer research, but many other interesting screening models such as differentiated iPSC cells, primary tissues, and difficult to transduce cells have been challenging to approach with genome-wide CRISPR screens (7, 8, 50). The optimized CRISPRi and CRISPRa libraries described in this work provide a resource that make these models more tractable for genetic screens.

The optimizations developed here can be used to clone even more compact libraries, such as multi-guide Cas9 and Cas12a sgRNA constructs for CRISPRi and CRISPRa perturbations (51, 52). With these multi-guide libraries, it is conceivable to have a guide library containing a single element per gene. A 20,000-guide library at 50-fold cell coverage only requires 1 million cells which can be propagated in a single 100mm dish or multi-well plate. With automation this can enable large panels for drugs to be screened using a genome-wide guide library, something unconceivable with previous library cloning techniques.

## Supporting information

Detailed cloning protocol

Primers for NGS library prep

Pilot cloning guide counts table

CRISPR library guide sequences

Guide plasmid library counts table

1000x vs 100x screen guide counts table

Transduction experiment guide counts table

Dasatinib screen guide counts table

## AVAILABILITY

ScreenProcessing is an open source bioinformatics pipeline available in the LGR’s GitHub repository (https://github.com/ucsf-lgr/ScreenProcessing).

MAGeCK-Vispr is an open source pipeline developed by Wei Lei and Han Xu from the Dr.Xiaole Shirley Liu laboratory and is available on SOURCEFORCE (https://sourceforge.net/p/mageck/wiki/Home/).

ClusterProfiler is an open source enrichment tool analysis R package available on Bioconductor (doi: 10.18129/B9.bioc.clusterProfiler).

The Database for Annotation Visualization and Integrated Discovery (DAVID) is an open source functional annotation tool that is available online (https://david.ncifcrf.gov/home.jsp).

## SUPPLEMENTARY DATA

Supplementary Data are available online.

## ACKNOWLEDGEMENT

We would like to thank Luke A. Gilbert for providing helpful feedback and comments on the manuscript. We also thank Karl Mader, Samira Yitiz, Isabella Turcinovic, Brandon Kwan-Leong and the LGR staff for technical support.

## TABLE AND FIGURE LEGENDS

Figures were created with BioRender.com.

