## Supplemental Data for "Optimized CRISPR guide RNA library cloning reduces skew and enables more compact genetic screens"

### Sup Fig. S1

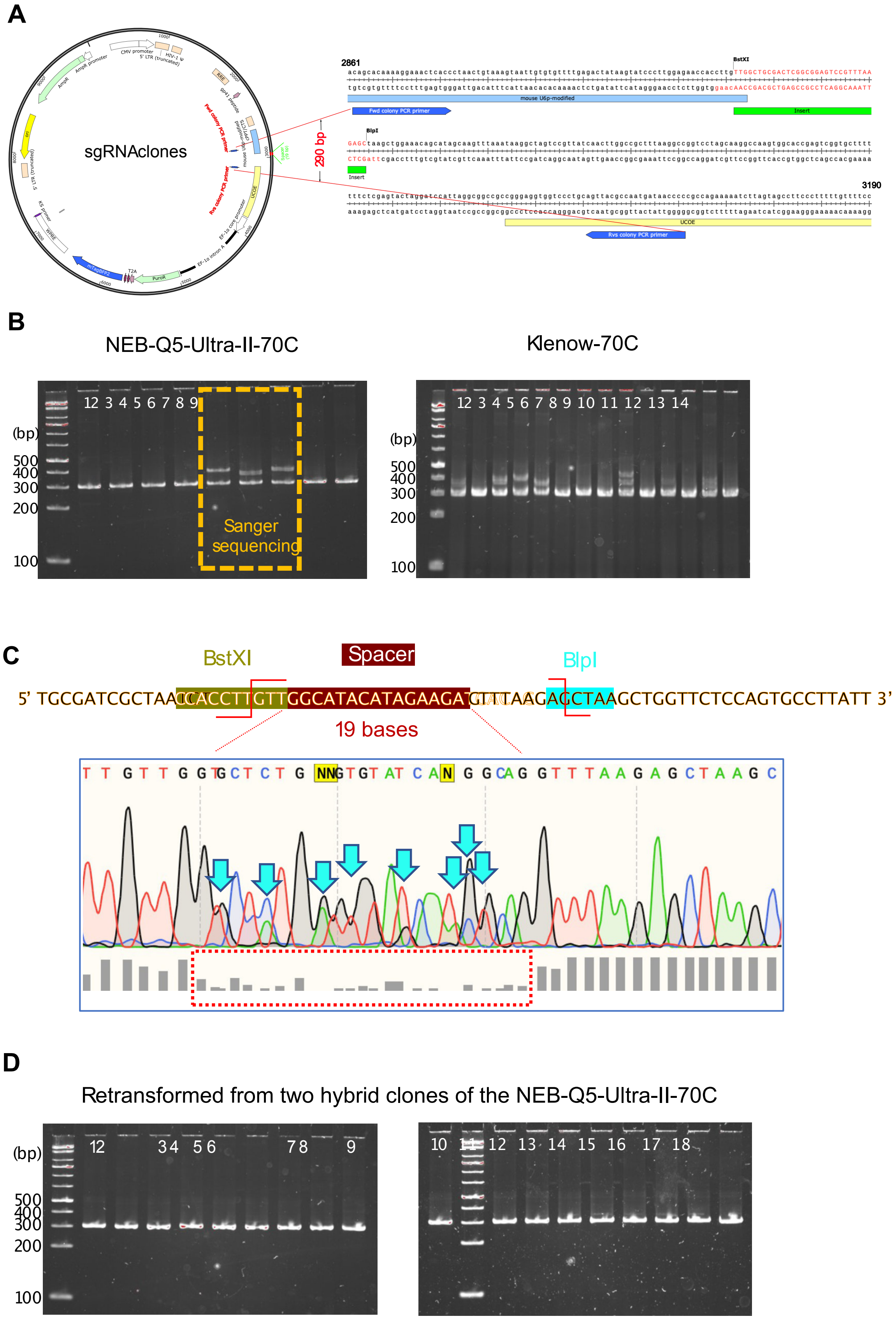
A
sgRNAclones
2861
BstXI
BlpI
3190
290 bp
Fwd colony PCR primer
Rvs colony PCR primer
Insert
UCOE
B
NEB-Q5-Ultra-II-70C
Klenow-70C
(bp)
500
400
300
200
100
Sanger sequencing
C
BstXI
Spacer
BlpI
5' TGCGATCGCTAACCACCTTGTTGGCATACATAGAAGATGTTAAGAGCTAAGCTGGTTCTCCAGTGCCTTATT 3'
19 bases
D
Retransformed from two hybrid clones of the NEB-Q5-Ultra-II-70C

Supplementary Figure S1. (A) Diagram of the cloning strategy and colony PCR primer design. (B) Colony PCR of sgRNA clones and gel electrophoresis. (C) Representative Sanger sequencing trace of plasmids from hybrid clones (D) Plasmids from two different hybrid clones were retransformed. Colony PCR was performed, and hybrid clones were not detected.

#### Sup Fig. S2

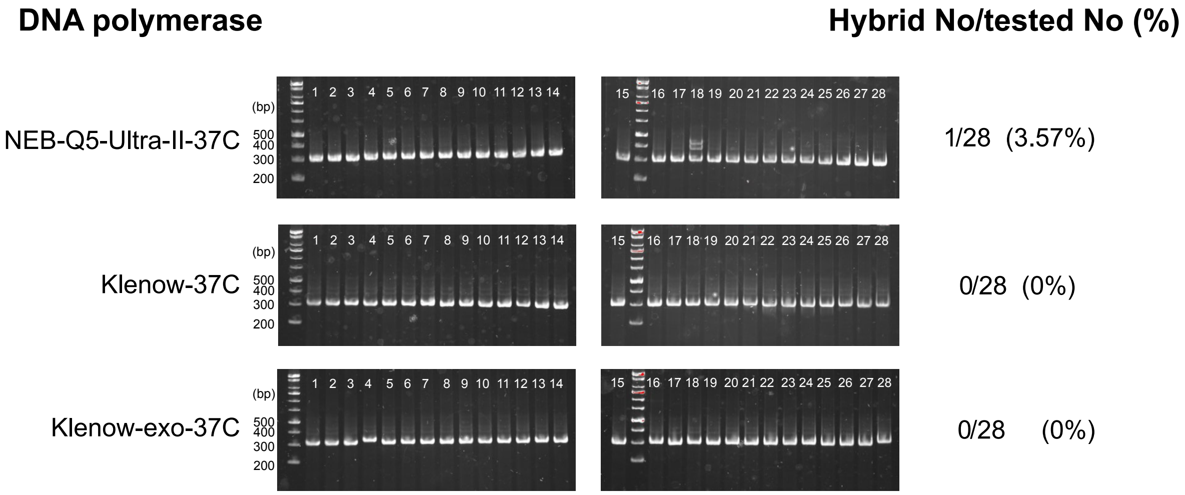


Supplementary Figure S2. Lowering the elution temperature for the insert gel extraction reduces the formation of hybrid clones, regardless of the insert preparation polymerase used.

#### Sup Fig. S3

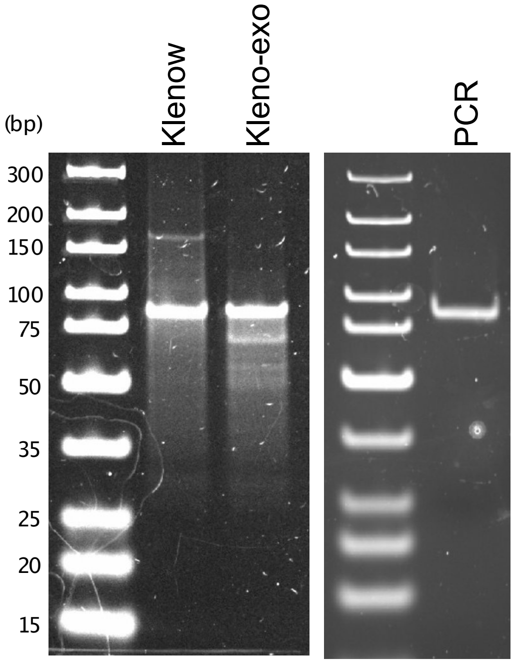


Supplementary Figure S3. Synthesis of double-stranded sgRNA with the three different DNA polymerases (Klenow, Klenow exo-, and NEB Q5 Ultra II) and gel electrophoresis indicates less by-products with PCR.

#### Sup Fig. S4

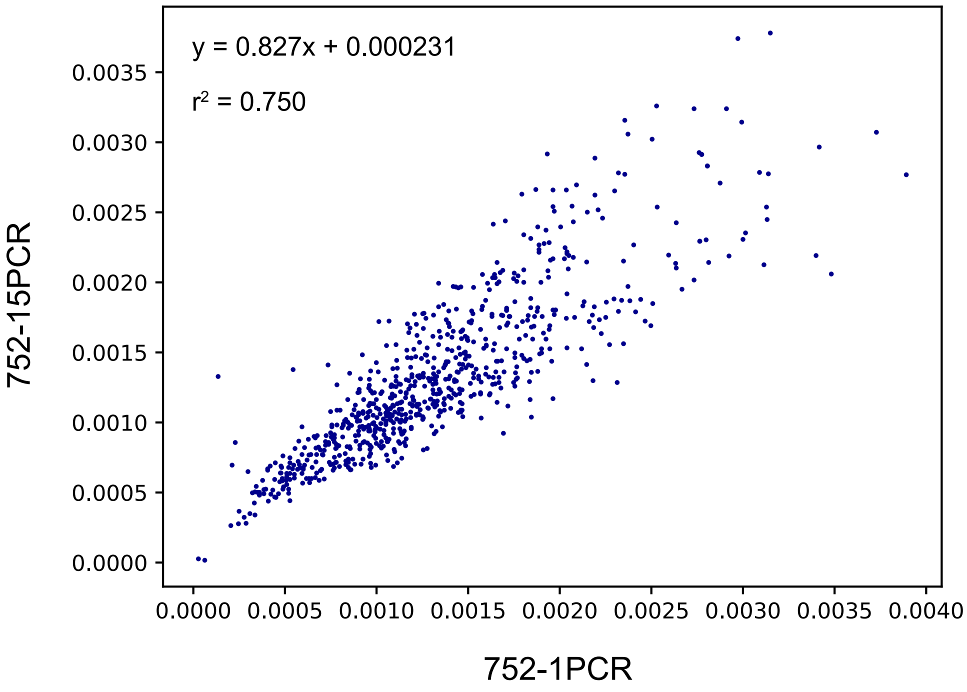


Supplementary Figure S4. Additional PCR cycles introduce minimal bias in sgRNA libraries. An oligo pool containing 752 guides was amplified by one or 15 cycles of PCR, digested, and cloned into a guide vector. A pairwise comparison of sequence counts of guides in each library is depicted.

**Sup Fig. S5**

**A**

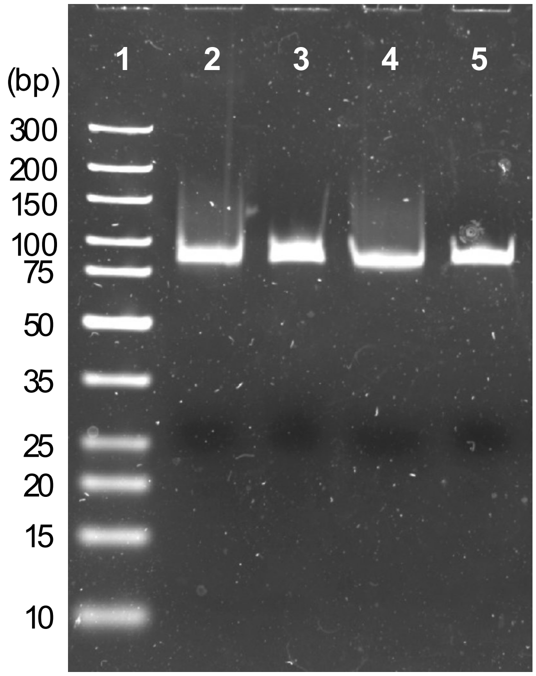
1
2
3
4
5
(bp)
300
200
150
100
75
50
35
25
20
15
10

**B**

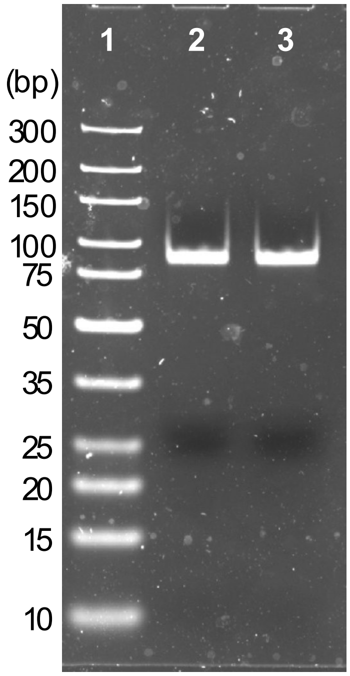
1
2
3
(bp)
300
200
150
100
75
50
35
25
20
15
10

Supplementary Figure S5. Optimization of PCR cycles and input template concentration reduces overamplification products. (A) Lane 1: DNA ladder, Lanes 2 and 4: 200pM template. Lanes 3 and 5: 10pM template. Lanes 2-5: 13 cycles PCR.  (B) Lane 1: DNA ladder. Lane 2: 200pM template, Lane 3: 600pM template. Lanes 2 and 3: 8 cycles PCR.

### Sup Fig. S6

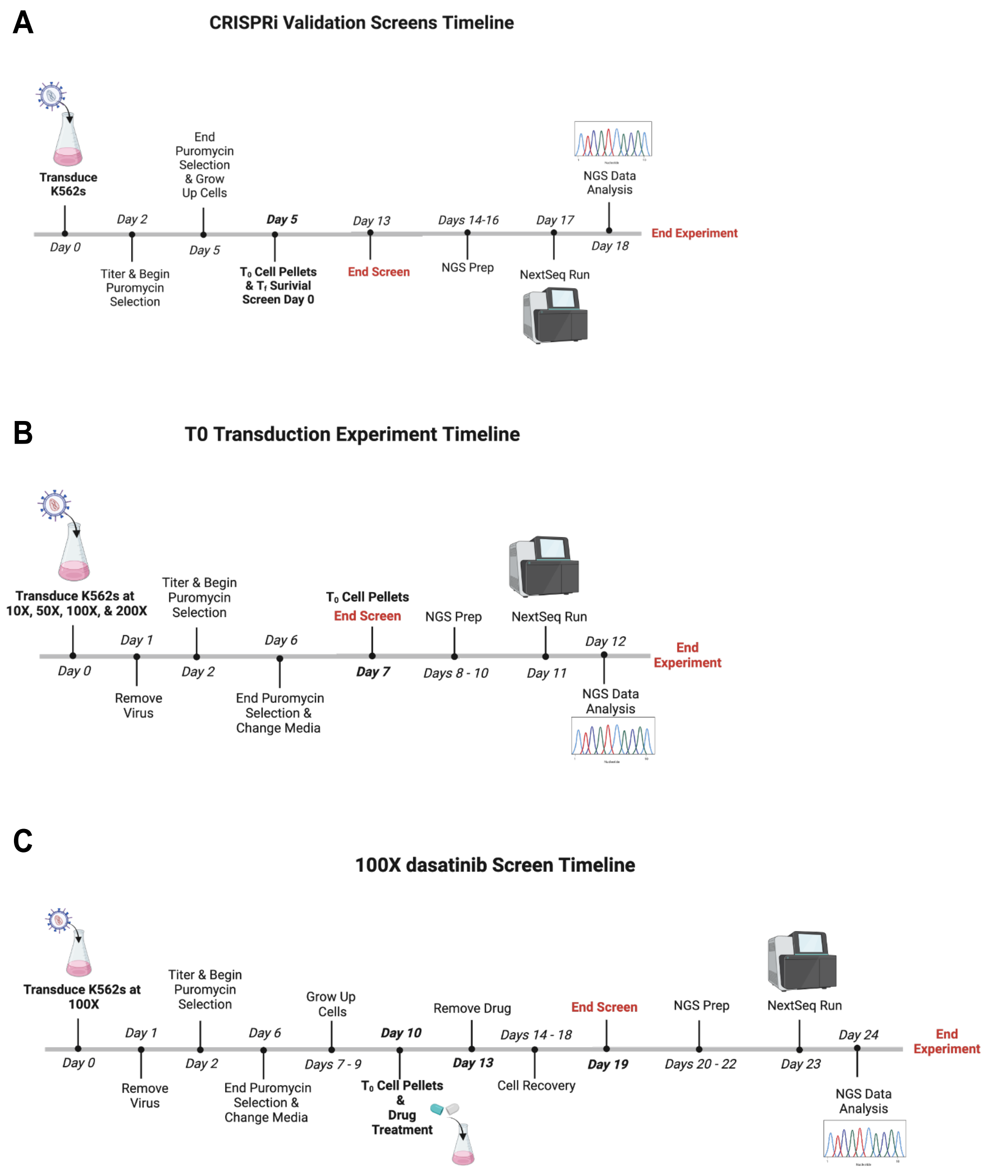


Supplementary Figure S6. Schematic of the screening timelines for the CRISPRi V2 validation screen (Figure 3), the transduction titration experiment (Figure 4), and the dasatinib survival screen (Figure 5). Timelines include the transduction duration, virus titer determination, puromycin selection length, cell pellet collections, and next-generation sequencing (NGS) sample preparation for each experiment. (A) CRISPRi V2 LGR 1000 and 100-fold cell coverage survival screen timeline. (B) CRISPRi V2 LGR verses legacy transduction titration experiment timeline. (C) LGR verses legacy 100-fold cell coverage dasatinib survival screen timeline.

#### Sup Fig. S7

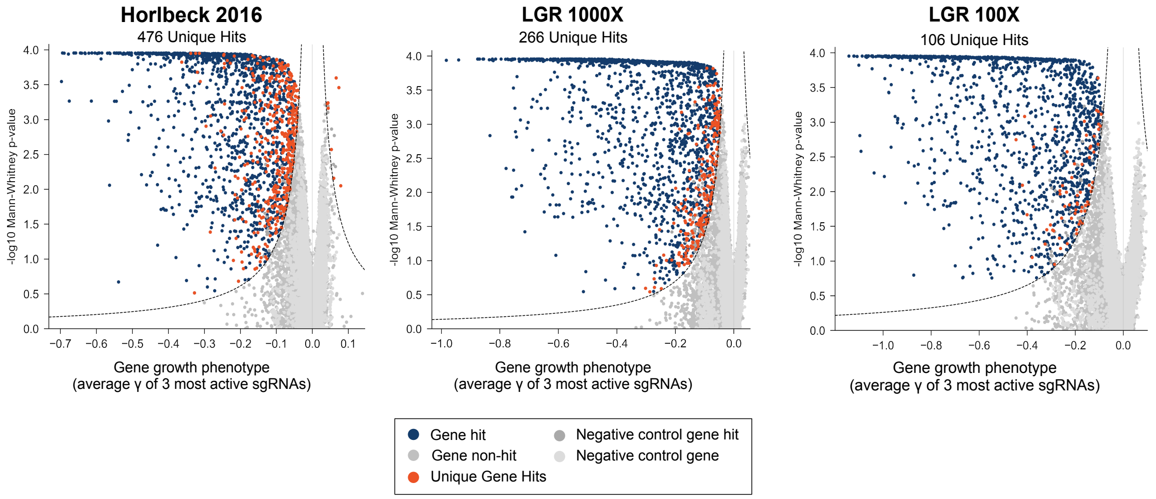


Supplementary Figure S7. Volcano plots of all three survival screens (Horlbeck et al. 2016, LGR 1000-fold cell coverage, and LGR 100-fold cell coverage). Unique gene hits identified for each library are orange and gene hits that are shared among the libraries are in blue. Unique gene hits for each library are near the cutoff (dotted lines) which are populated by weaker and/or less significant hits.

### Sup Fig. S8

A

| Sample | Total sgRNAs Drop Out | % Total Guides | % of Poly G Drop Outs |
|---|---|---|---|
| LGR_10X_1 | 765 | 0.732 | 8.627 |
| LGR_10X_2 | 553 | 0.529 | 11.754 |
| Legacy_10X_1 | 7409 | 7.088 | 1.039 |
| Legacy_10X_2 | 6141 | 5.875 | 1.238 |
| LGR_50X_1 | 72 | 0.069 | 90.278 |
| LGR_50X_2 | 71 | 0.068 | 90.141 |
| Legacy_50X_1 | 1092 | 1.045 | 5.861 |
| Legacy_50X_2 | 1056 | 1.010 | 6.155 |
| LGR_100X_1 | 69 | 0.066 | 94.203 |
| LGR_100X_2 | 71 | 0.068 | 92.958 |
| Legacy_100X_1 | 950 | 0.909 | 6.842 |
| Legacy_100X_2 | 841 | 0.805 | 7.729 |
| LGR_200X_1 | 71 | 0.068 | 88.732 |
| LGR_200X_2 | 71 | 0.068 | 88.732 |
| Legacy_200X_1 | 844 | 0.807 | 7.464 |
| Legacy_200X_2 | 860 | 0.823 | 7.093 |
| Plasmid LGR NextSeq | 59 | 0.056 | 93.220 |
| Plasmid LGR HiSeq | 2 | 0.002 | 0.000 |
| Plasmid Legacy NextSeq | 770 | 0.737 | 14.545 |

B

| LGR CRISPRi Plasmid HiSeq | | LGR CRISPRi Plasmid NextSeq | |
|---|---|---|---|
| Skew : 1.92 | | Skew : 1.99 | |
| **Frequency of Drop Out** | **First 5 Bases of sgRNA** | **Frequency of Drop Out** | **First 5 Bases of sgRNA** |
| 1 | GTCAC | 55 | GGGGG |
| 1 | GCCCG | 1 | GTCAC |
| | | 1 | GGGGA |
| | | 1 | GGCTC |

Supplementary Figure S8. The majority of dropouts identified in the CRISPRi V2 LGR library (plasmid and K562 samples) was due to the Illumina NextSeq 550 sequencing artifact. The NextSeq 550 is a two-color sequencer that requires signal in the first few cycles to find clusters on the flow cell. Since G bases are dark and lack signal, sgRNAs that begin with a polyG are not detected. (A) In a 103,073 element library, the majority of guides that drop out of the LGR library samples at higher fold cell coverage (50, 100, and 200) are sgRNAs that begin with polyG sequences. Whereas the legacy library samples at the same cell coverages experienced dropouts due to the library quality. (B) Sequence composition analysis of the CRISPRi V2 LGR plasmid library determined that most sgRNAs dropping out began with the polyG sequence in the NextSeq 550 run. When the same library was sequence on the four-color HiSeq 4000, much fewer guides dropped out and a lower 90/10 skew ratio was achieved.

#### Sup Fig. S9

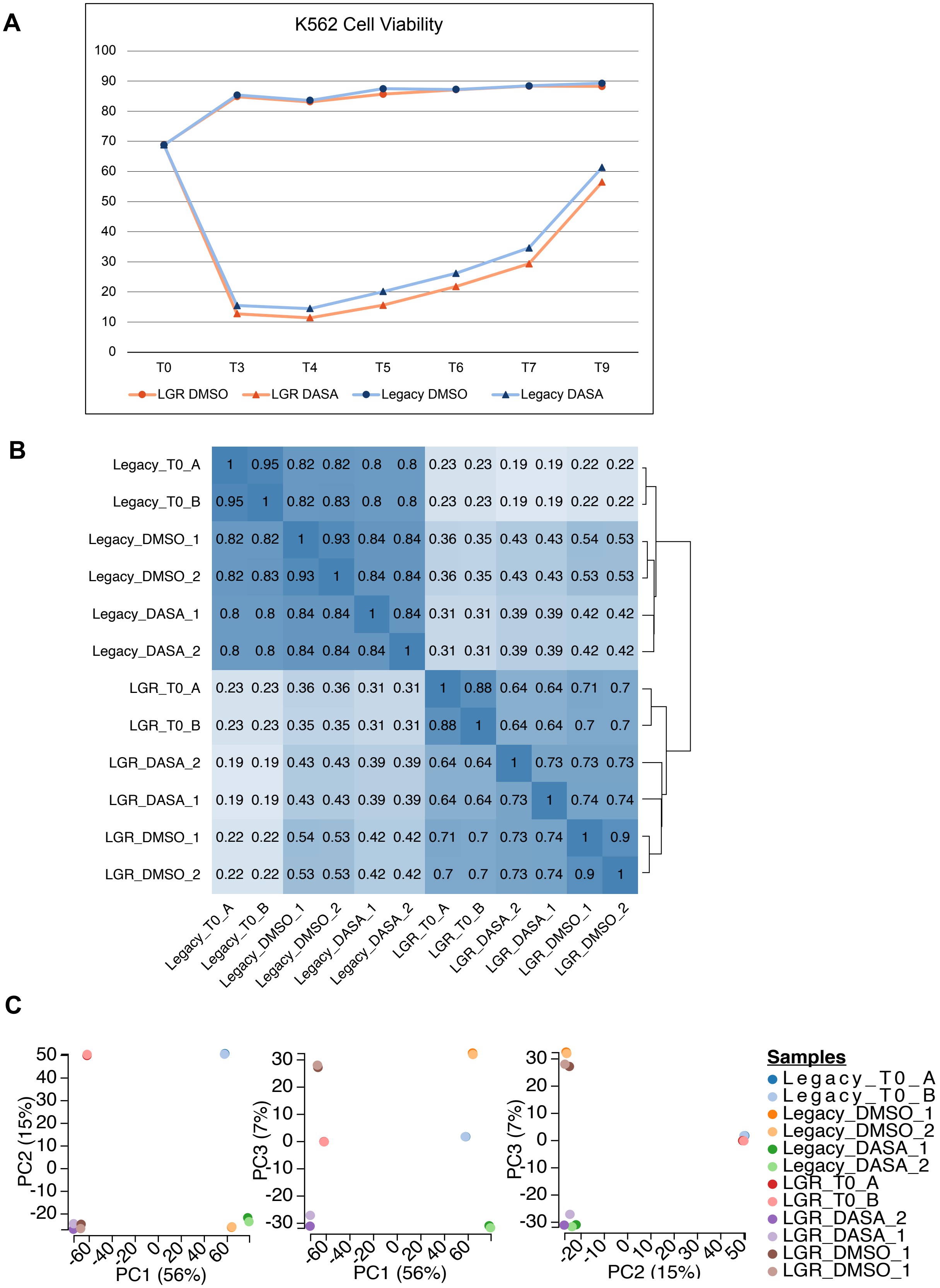
A
K562 Cell Viability
100
90
80
70
60
50
40
30
20
10
0
T0
T3
T4
T5
T6
T7
T9
LGR DMSO
LGR DASA
Legacy DMSO
Legacy DASA
B
Legacy_T0_A 1 0.95 0.82 0.82 0.8 0.8 0.23 0.23 0.19 0.19 0.22 0.22
Legacy_T0_B 0.95 1 0.82 0.83 0.8 0.8 0.23 0.23 0.19 0.19 0.22 0.22
Legacy_DMSO_1 0.82 0.82 1 0.93 0.84 0.84 0.36 0.35 0.43 0.43 0.54 0.53
Legacy_DMSO_2 0.82 0.83 0.93 1 0.84 0.84 0.36 0.35 0.43 0.43 0.53 0.53
Legacy_DASA_1 0.8 0.8 0.84 0.84 1 0.84 0.31 0.31 0.39 0.39 0.42 0.42
Legacy_DASA_2 0.8 0.8 0.84 0.84 0.84 1 0.31 0.31 0.39 0.39 0.42 0.42
LGR_T0_A 0.23 0.23 0.36 0.36 0.31 0.31 1 0.88 0.64 0.64 0.71 0.7
LGR_T0_B 0.23 0.23 0.35 0.35 0.31 0.31 0.88 1 0.64 0.64 0.7 0.7
LGR_DASA_2 0.19 0.19 0.43 0.43 0.39 0.39 0.64 0.64 1 0.73 0.73 0.73
LGR_DASA_1 0.19 0.19 0.43 0.43 0.39 0.39 0.64 0.64 0.73 1 0.74 0.74
LGR_DMSO_1 0.22 0.22 0.54 0.53 0.42 0.42 0.71 0.7 0.73 0.74 1 0.9
LGR_DMSO_2 0.22 0.22 0.53 0.53 0.42 0.42 0.7 0.7 0.73 0.74 0.9 1
Legacy_T0_A
Legacy_T0_B
Legacy_DMSO_1
Legacy_DMSO_2
Legacy_DASA_1
Legacy_DASA_2
LGR_T0_A
LGR_T0_B
LGR_DASA_2
LGR_DASA_1
LGR_DMSO_1
LGR_DMSO_2
C
PC2 (15%)
PC1 (56%)
PC3 (7%)
PC1 (56%)
PC3 (7%)
PC2 (15%)
Samples
Legacy_T0_A
Legacy_T0_B
Legacy_DMSO_1
Legacy_DMSO_2
Legacy_DASA_1
Legacy_DASA_2
LGR_T0_A
LGR_T0_B
LGR_DASA_2
LGR_DASA_1
LGR_DMSO_1
LGR_DMSO_1

Supplementary Figure S9. Dasatinib survival screen quality control measurements for the LGR and legacy library samples. (A) K562 cell viability graph for the average LGR and legacy dasatinib treated (DASA) and control (DMSO) samples over the 10 days of the experiment. Dasatinib treatment was for 72 hours (T0 - T3) and recovery was for 5 days (T3 – T9). (B) Hierarchical clustering of the LGR and legacy library samples. (C) Principal Component Analysis (PCA) plot of the LGR and legacy library samples.

### Sup Fig. S10

#### Top 12 Barplots of GO & KEGG annotations
#### FDR 0.001

##### Legacy
##### Positive Gene Hits

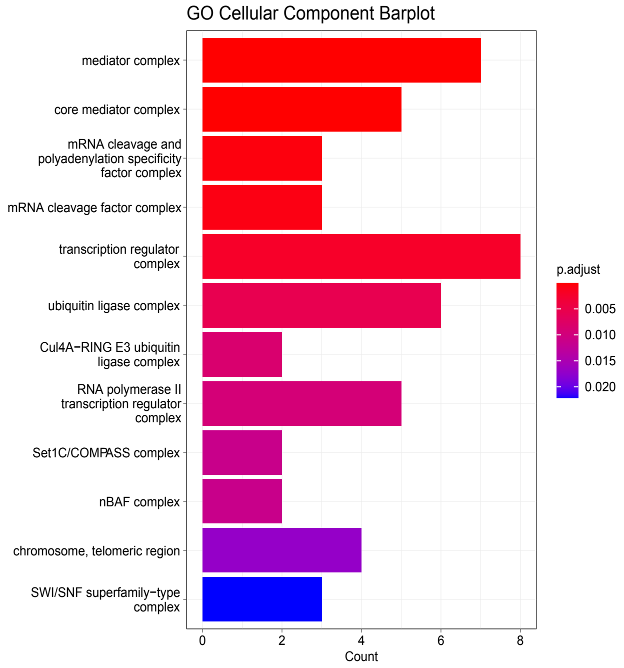


##### LGR
##### Positive Gene Hits

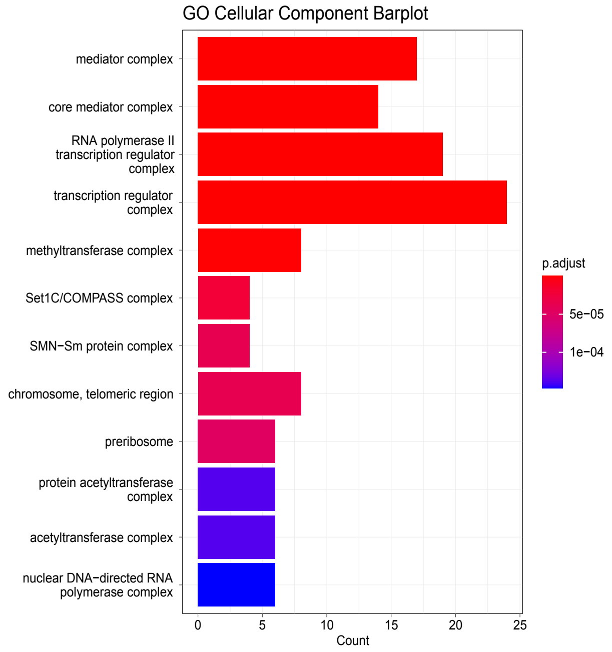


##### Legacy
##### Negative Gene Hits

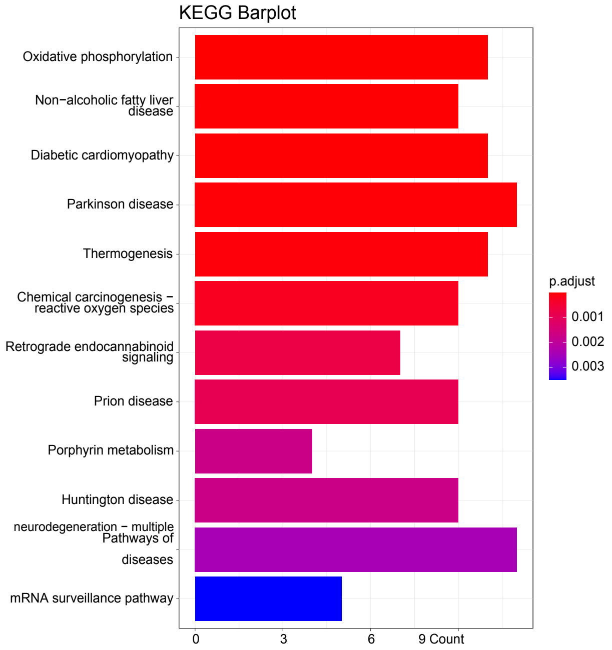


##### LGR
##### Negative Gene Hits

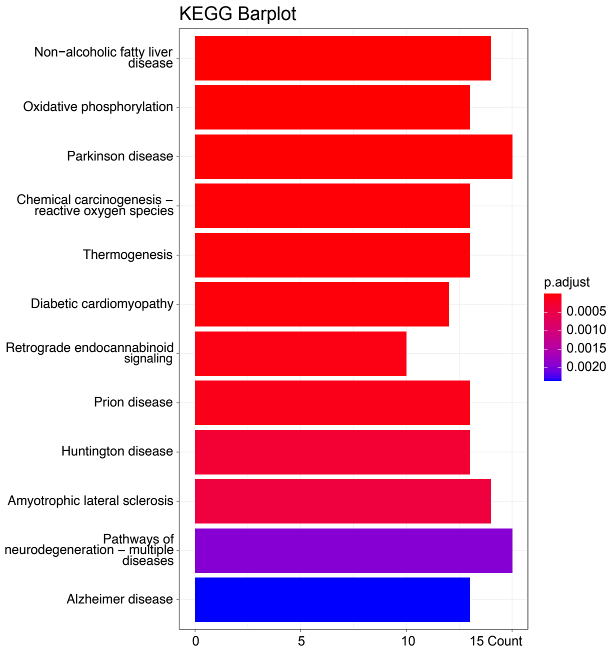

Supplementary Figure S10. Bar plots ranking the top 12 positive and negative gene hits for the 100-fold cell coverage dasatinib survival screen performed with the legacy (left) and LGR (right) libraries. Gene hits are ranked by p-value assigned by ClusterProfiler.

### Sup Fig. S11

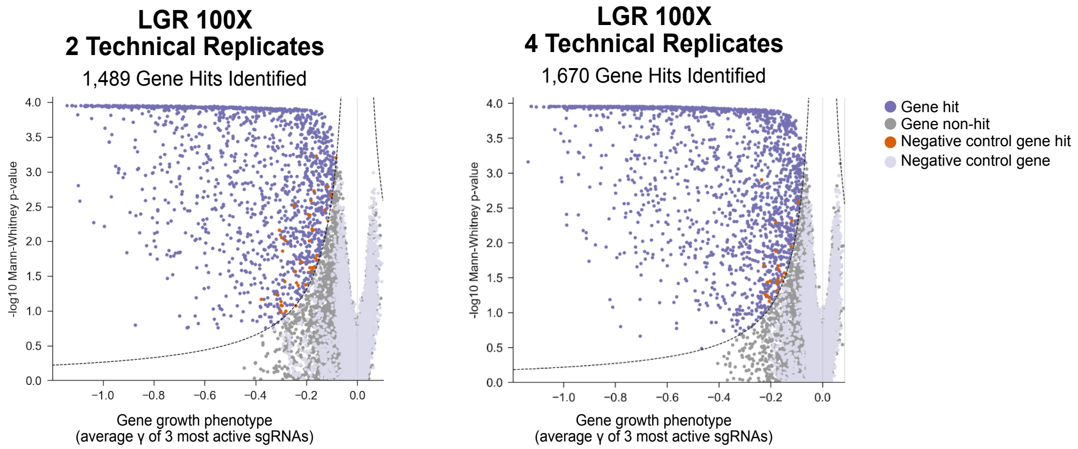


Supplementary Figure S11. The total number of essential genes identified in the LGR  100-fold cell screen increases as more technical replicates are processed.

#### Sup Fig. S12

| Screening Coverage | Number of Infected Cells for 100,000 Guide Library | Number of Cells to Transduce with a 25 % Infection | Suspension Cell Line Culture Volume for 1 Sample *(with 3 technical replicates at ~500,000 cells/mL)* | Suspension Cell Line Shaker Flask Size for 1 Sample | Adherent Cell Line Flask Size for 1 Sample |
|---|---|---|---|---|---|
| 10X | 1 million | 4 million | 6 mL *(30 mL minimum 125mL culture volume)* | 125 mL | T25 |
| 50X | 5 million | 20 million | 30 mL | 125 mL | T75 |
| 100X | 10 million | 40 million | 60 mL | 125 mL | T175 |
| 200X | 20 million | 80 million | 120 mL | 250 mL | T175 |
| 500X | 50 million | 200 million | 300 mL | 500 mL | Two T225 |
| 1000X | 100 million | 400 million | 600 mL | 1 L | Four T225 |

Supplementary Figure S12. A table demonstrating the quantity of materials necessary to perform genome-wide screens at increasing coverages (10 to 1000-fold cell coverage). Labware recommendations were acquired from ThermoFisher.
