## Supplementary material for "Optimized CRISPR guide RNA library cloning reduces skew and enables more compact genetic screens": Detailed cloning protocol

### LGR CRISPR Guide Cloning Protocol

#### Introduction

This protocol is used to clone guide RNAs into the LGR CRISPR guide vector.

#### Steps

1. Generate oligo sequences.
2. Order oligo pool.
3. PCR amplification of oligo pools.
4. BlpI/BstXI restriction digest and purification of oligo pool PCR product (guide insert).
5. BlpI/BstXI restriction digest of guide vector.
6. Ligation of insert into vector.
7. Transformation of ligation product.
8. Plasmid purification of cloned library.

#### Materials

NEBNext Ultra II Q5 Master Mix (NEB #M0544S)
NEB BlpI (NEB #R0585S)
NEB BstXI (NEB #R0113S)
NEB buffers: 10x buffer 2.1 and 3.1
High Sensitivity dsDNA Quantitation Kit (Biotium #31066)
NEB T4 DNA Ligase (NEB #M0202S)
NEB 5-alpha Competent *E. coli* (High Efficiency) (NEB #C2987)
Thermo Fisher MegaX DH10B Electrocomp™ Cells (ThermoFsher #C640003)
Qiagen MinElute PCR Purification Kit (Qiagen #28004)
Qiagen QIAquick Nucleotide Removal Kit (Qiagen #28306)
Qiagen QIAprep Spin Miniprep Kit (Qiagen #27104)
NucleoSpin Gel and PCR Clean-up Kit (Macherey-Nagel #740609.50)
TriTrack DNA Loading Dye (6X) (ThermoFisher #R1161)
Low Melt Agarose (Bio-Rad #161311)
Mini-Protean Tetra Vertical Electrophoresis Cell (Bio-Rad #165-8000)
10X TBE buffer
GeneRuler Ultra Low Range DNA Ladder (ThermoFisher # SM1211)
GeneRuler 1 kb Plus DNA Ladder (ThermoFisher #SM1334)
30% Acrylamide/Bis Solution (29:1) (Bio-Rad #1610156)
10% Ammonium Persulfate
TEMED
Ammonium acetate (pH8.0)
Magnesium acetate tetrahydrate
Sodium acetate (3M, pH 5.2)
EDTA, pH 8.0
Low EDTA TE (1X), pH 8.0 (VWR #10128-588)
1M Tris HCl, pH 7.9
500 mM NaCl
0.5 mL Nonstick, RNAse-free Microfuge Tubes (Ambion #AM12350)
1.5 mL Nonstick, RNAse-free Microfuge Tubes (Ambion #AM12450)

18.5 gauge needle
GelGreen Nucleic Acid Stain (Biotium #41005)
UV Fluorescent Ruler
Costar Spin-X Centrifuge Tube Filters (Sigma #CLS8160)
GlycoBlue Coprecipitant (ThermoFisher #AM9515)
ZymoPURE II Plasmid Maxiprep Kit (Zymo Research # D4203)

**Step 1. Generating oligo sequences**

The dual guide vector is designed to accommodate 20 nt guide sequences ("protospacers"), the first base of which must be a "G" due to the preference of the U6 promoter. The first base of each spacer should be changed to a G. Each oligo is 84 nucleotides (nt) long and have the following structure.

5'- PCR adapter - ccaccttgttg - protospacer - gtttaagagctaagctg - PCR adapter-3'

The 5'- and 3'- PCR adapters serve as handles for PCR amplification. If you want to generate multiple subpools from a single oligo pool, design each subpool to have different 5'- and 3'- PCR adapter pairs. (Note: It is important for the subpool representations to be approximately balanced, as a lowly represented subpools will be difficult to PCR amplify). The full list of 5'- and 3'- PCR adapters can be found at the end of the protocol. To avoid cross contamination, you should order subsequent oligo pools with new PCR adapter sequences.

**example single guide oligo:**

CGTAGCGTTTTGTACACGccaccttgttgGAAAATGAGGTGCATAAGGAgtttaagagctaagctgCACGCTCAAAAAGCGTAC

In addition to appending the 5'- and 3'- PCR adapters to the oligo sequences, you will also need to order the corresponding primers to amplify the oligo pool.

**Step 2. Ordering oligos**

We have been sourcing our oligos from Agilent (Microarray Oligo Library Synthesis) and IDT (oPools).

When purchasing oligos, order both forward and reverse complement sequences to minimize sequence-specific biases. Agilent has a few different scales of synthesis to deliver ~10 pmol oligo per pool, regardless of the number of oligos you order. To do this, they make extra copies of the oligos once the total oligo number drops below certain levels. Note that these synthesis tiers don't match pricing tiers (see table below). For our library, we ordered several oligo pools that didn't exceed 55,000 elements each

IDT oPools are synthesized individually at small and then pooled. To cut down on cost, the synthesis reactions are not quantified. Instead, algorithm created from empirical data estimates the amount of oligo produced and then depending on the pool size 1, 10, or 50 pmol of each oligo are combined lyophilized.

Depending on the size of your oligo pool and cost of IDT and Agilent synthesis, it might be more cost effective to use an Agilent synthesis once you hit 500 or 1000 guides.

**Step 3. PCR amplification of oligo pools (Day 1)**
*Setup of this PCR reaction should be completed in a PCR-free area (see appendix) to avoid contamination with other PCR products, plasmids, and libraries.

Depending on the amount of oligo delivered, re-suspend your oligo pool in low EDTA TE to produce a 0.1 pmol/µL stock. Store this stock in a -20˚C in a PCR-free area.

Dilute an aliquot of the stock in low EDTA TE to produce a 0.01 pmol/µL oligo pool.

**Set up PCR reaction**

| | standard reaction |
|---|---|
| NEBNext Ultra II Q5 Master Mix | 50 µL |
| 50 uM Premixed F- and R-primer (final 500 nM) | 1 µL |
| Template (0.01 pmol/µL, final 200 pM) | 2 µL |
| H2O | 47 µL |
| **Total** | **100 µL** |

**Run PCR reaction**
Calculate the appropriate annealing temps for each primer pair using the NEB Tm calculator for Q5: https://tmcalculator.neb.com/#!/main

| | temperature | duration |
|---|---|---|
| Initial Denaturation | 98°C | 30 seconds |
| Denaturation | 98°C | 10 seconds |
| Annealing | variable | 20 seconds |
| Extension | 72°C | 15 seconds |
| Final Extension | 72°C | 2 minutes |
| Hold | 4°C | ∞ |

In our hands, 7 cycles of PCR generated enough template to clone our libraries twice. You may want to increase the number of cycles if you don't get enough material.

1. Clean PCR products using the Qiagen PCR MinElute Kit, eluting in 20 µL Buffer EB.
2. Run PCR product on TapeStation to check product size. The expected product size is 84 bp. If there are non-specific bands, do not proceed to the next step because non-specific inserts may be cloned into your pool.
3. Quantify PCR product concentration with the fluorometric Biotium or Qubit High sensitivity assay. A NanoDrop will not provide accurate quantitation at lower concentrations.

**Step 4. BlpI/BstXI restriction digest and purification of guide insert.**
Sequentially digest the PCR product with BlpI/BstXI to produce the appropriate sticky ends for ligation into the guide vector.

Prior to beginning digestions, prepare 1.5L of 0.5X TBE and store at 4˚C. This will be used during electrophoresis.

**Oligo Pool BlpI digestion (43 µL reaction)**

| | standard |
|---|---|
| oligo pool PCR product | 50 ng |
| NEBuffer 2.1 | 4.3 µL |
| BlpI (NEB) | 4 µL |
| H2O | variable |
| **total** | **43 µL** |

1. Mix well and spin down very briefly
2. Incubate in the PCR machine (37˚C, 15 min; Lid temperature: 47˚C)
3. Immediately clean up digestion using the Qiagen Nucleotide Removal Kit. Note we increased the drying time to ensure removal of all of the wash buffer:
    a. Mix 430 µL of PNI Buffer with your sample and apply it to the column.
    b. Spin at 6000rpm, 1 min
    c. Add 750 µL Buffer PE, 6000rpm
    d. Dry for 1.5 min at 13,000 rpm
    e. Elute with 53 µL of Buffer EB

**Oligo Pool BstXI digestion (60 µL reaction)**

| | standard |
|---|---|
| BlpI reaction product | 51 µL |
| NEBuffer 3.1 | 6 µL |
| BstXI (NEB) | 3 µL |
| **total** | **60 µL** |

1. Mix well and spin down very briefly
2. Incubate in the PCR machine (37˚C, 15 min; Lid temperature: 47˚C)
3. After 15 min digestion, immediately add 6x loading dye, mix well, and keep on ice.

**Electrophoresis and purification of guide insert**

Digestion of a PCR product is expected to produce 26, 33, and 25 bp fragments, use a 10% polyacrylamide gel to properly resolve the desired 33 bp insert from the two smaller fragments.

**Pour polyacrylamide gel**

| | 10% gel |
|---|---|
| 30% acrylamide | 3.96 mL |
| 10X TBE | 1.2 mL |
| 10% APS | 120 µL |
| H2O | 6.708 mL |
| TEMED | 12 µL |

**Run polyacrylamide gel**

Due to the short length of the digested insert, it is important to maintain guide stability and minimize denaturation in order to maintain uniform guide representation in the library. To achieve this, we run the gel under low voltage at 4˚C. This is also the reason the TBE buffer is pre-chilled.

1. Find a styrofoam box that is large enough to fit the gel box, with 1-2 inch margins between the gel box and the styrofoam box.
2. Place the gel box inside the styrofoam, filling the bottom and side margins with ice (see example below).
3. Place gel in gel box and fill with prepared 0.5x TBE pre-chilled to 4˚C.
4. Load the gel with the reaction product from step (b) in two wells.
5. Run gel at 80V until the yellow dye (~15 bp) has traveled at the end of the gel. This usually takes 2-3h. The gel is run out as far as possible to separate the insert from the 5' and 3' adapters, as well as from any potentially undigested fragments.

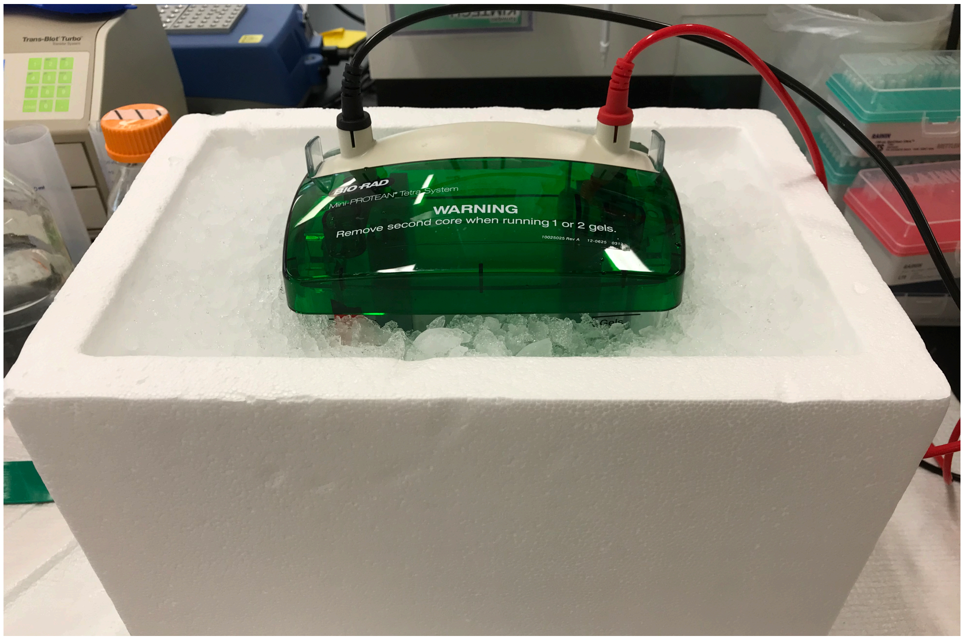


**Prepare cold 1x Acrylamide Gel Elution Buffer**

| | stock conc. | final conc. | volume added |
|---|---|---|---|
| ammonium acetate | 7.5M | 500 mM | 71.43 µL |
| magnesium acetate tetrahydrate | 1M | 10 mM | 10.0 µL |
| EDTA, pH 8.0 | 0.5M | 1 mM | 2.0 µL |
| H2O | -- | -- | 916.57 µL |
| **total** | | | **1 mL** |

**Extract and elute guide insert**

1. The gel slice containing the insert will be disrupted using a set of nested centrifuge tubes. To prepare the tubes, puncture the bottom of a 0.5 mL nonstick microtube with an 18.5 gauge needle and place inside a 1.5 mL nonstick microtube.

2. Next, stain the acrylamide gel in a small tray containing 50 mL cold 0.5x TBE from the electrophoresis rig and 5 µL 10,000x GelGreen stain. Cover with foil to protect the dye from light and gently incubate on a rocker for 2-3 minutes at room temperature.
3. Using a gel imaging station and the UV fluorescent ruler, cut out the band of interest, located around 35 bp, and place it into two pre-punctured 0.5 mL microtubes. Set the 0.5 mL tube inside a 1.5 mL microtube.
4. Spin at 21,130 rcf for 2 minutes to force the gel through the hole. This will break the gel slice into smaller fragments to aid diffusion. If the gel has not fully passed though, spin again.
5. Once all of the gel has passed through, add 300 µL cold elution buffer to each sample (no need to mix). Incubate without any shaking at 4˚C overnight to allow time for insert to diffuse out of the gel.

**Extract and elute guide insert (Day 2)**

1. Use a cut P1000 tip to transfer the gel slurry to 2 Costar Spin-X columns. You will have two tubes per sample.
2. Centrifuge Spin-X columns at 16,873g for 3 minutes.
3. Check for recovery of the full elution buffer volume; if recovery volume is lower than expected, turn the filter 180 degrees and re-spin to recover remaining elution.
4. Add, in the following order, on ice:
5. 3 µL glycoblue coprecipitant
6. 35 µL 3M sodium acetate
7. Add 875 µL ice cold EtOH.
8. Mix gently by flicking the tube and incubate at -20˚C overnight to precipitate the insert.

**Extract and elute guide insert (Day 3)**

1. Spin the tube for 30 minutes at 20,000g at 4˚C to pellet the DNA.
2. Remove supernatant.
3. Wash pellet twice with 800 µL ice cold 80% EtOH (20,000g at 4˚C for 30 min).
4. Spin for 1 minute at 20,000g at 4˚C.
5. Remove residual EtOH, let pellet air dry.
6. Once pellet is dry, add <u>10 µL cold 1x T4 ligase buffer per tube (20 µL total for each insert preparation).</u>
7. Allow pellet to re-hydrate on ice and pipette to re-suspend very gently.
8. Measure the concentration using a high sensitivity Biotium or Qubit high sensitivity assay

**<u>Step 5. BlpI/BstXI restriction digest of guide vector.</u>**

Sequentially digest the dual guide vector purified from the mini-prep with BlpI/BstXI to produce the appropriate sticky ends for ligation with the guide insert prepared in Step 4. This can be prepared ahead of time and stored in the freezer.

**Guide Vector BlpI digestion**

If you are cloning a genome-wide library, make two reactions

| | volume (µL) |
|---|---|
| Vector (400 ng) | 400 ng |
| BlpI (NEB) | 1 |
| 10x NEBuffer 2.1 | 1.25 |
| H2O | variable |
| **total volume** | **12.5** |

1. Mix well and spin down very briefly
2. Incubate in the PCR machine (37˚C, 15 min)
3. Proceed immediately to BstXI digestion.

**Guide Vector BstXI digestion**

| | volume (µL) |
|---|---|
| BlpI reaction product | 12.5 |
| BstXI (NEB) | 1 |
| 10x NEBuffer 2.1 | 1.25 |
| Tris-HCl (1M, pH 7.9) | 1 |
| NaCl (500 mM) | 5 |
| H2O | 4.25 |
| **total volume** | **25** |

1. Mix well and spin down very briefly
2. Incubate in the PCR machine (37˚C, 15 min)
3. After 15 min digestion, immediately add 6x loading dye, mix well, and keep on ice.

**Gel Purification**

1. Prepare a 1% low-melt agarose gel using 1x TAE and 1x GelGreen nucleic acid stain. Low-melt agarose is used so the vector can be extracted in more gentle conditions
2. After loading samples, run gel at 75V until you get good separation between cut and uncut vector.
3. The gel is very soft. Handle the gel very carefully. Excise linearized vector, transfer into 1.5 mL microtube.
4. Gel extract with the Macherey-Nagel NucleoSpin Gel and PCR Clean-up Kit, with the following adjustments to preserve the quality of the insert:
    a. Pre-warm elution buffer (Buffer NE) to 70˚C.
    b. After adding NT1 buffer, mix gently at RT on a rocker for at least 10 minutes (instead of 50˚C with vortexing).
    c. Spin down very briefly and check for any gel debris.
    d. If there are any visible pieces of gel remaining, place back on rocker for 5 minutes.
    e. Once there are no visible pieces of gel remaining, place on rocker for another 5 minutes to allow pieces too small to see by eye to dissolve.
    f. After the final 5 minutes, briefly spin down and proceed with the protocol.
    g. For DNA binding and washing steps, centrifuge at 6,500g but not 11,000g.
    h. Increase the centrifugation time of Step 4 (column drying after the 2nd wash) to 90 seconds.
    i. Elute the digested vector in 25 µL pre-warmed elution buffer to increase recovery.
    j. Once eluted, measure the product using the Qubit.
    k. The yield should be about 50-60%.
5. Proceed to ligation in the next section and store left over prepared vectors at -20˚C.

**Step 6. Ligation of guide insert into guide vector.**
Perform a ligation using 1:2 vector:insert ratio. The guide vector is 9678 bp while the guide insert is 33 bp. For large genome-wide libraries, two or three ligation reactions will be required to get enough clones.

Additionally, set up a 1:0 vector:insert reaction as a self-ligation control. This will allow you to determine the background rate.

| | Negative Control | Guide insert |
|---|---|---|
| digested dual guide vector | 50 ng | 50 ng |
| digested guide insert | 0 ng | 0.34 ng |
| 10x ligation buffer | 2 µL | 2 µL |
| T4 DNA ligase | 1 µL | 1 µL |
| H2O | variable | variable |
| **total volume** | **20 µL** | **20 µL** |

Incubate ligations at 16˚C overnight.

**Step 7. Transformation of ligation product. (Day 4)**
Transform the ligation product into competent *E. coli*. For cloning libraries with greater than 1000 elements, we recommend transforming into electrocompetent cells. For smaller libraries, transforming into chemically competent cells is sufficient.

**Chemically competent transformation**
For both the full ligation reaction and the self-ligation control, transform 2 µL of ligation product into 50 µL NEB 5-alpha Competent *E. coli* (High Efficiency) and plate about 1% of liquid culture onto one LB plate to know the percent of background. Full ligation reaction with single guide insert can be expected to produce about more than 20,000 colonies from transformation with 2 µL of ligation. We aim to achieve 60x+ transformation coverage of our libraries. Depending on the number of elements in your library, you may need to set up multiple transformations.

The transformation protocol below is based on the NEB "High Efficiency Transformation" protocol.

1. Thaw a tube of NEB 5-alpha Competent *E. coli* (High Efficiency) cells on ice for 10 minutes.
2. Add 2 µL of ligation reaction to 50µL *E. coli* in a 1.5 mL microtube. Carefully flick the tube 4-5 times to mix cells and DNA. Do not vortex.
3. Place the mixture on ice for 30 minutes. Do not mix.
4. Heat shock at exactly 42°C for exactly 30 seconds. Do not mix.
5. Place on ice for 5 minutes. Do not mix.
6. Pipette 1000 µL of room temperature SOC into the mixture.
7. Place at 37°C for 60 minutes. Shake vigorously
8. Spread onto two selection plates and incubate overnight at 37°C.

**Electroporation**

Combine two ligations and perform ethanol precipitation to remove salts prior to electroporation

| | |
|---|---|
| Ligation | 40 µL |
| H2O | 160 µL |
| Glycoblue | 2 µL |
| 3M NaOAc | 20 µL |
| EtOH | 660 µL |

Incubate overnight at -20˚C.

**(Day 5)**

1. Centrifuge at 20,000g for 30 min at 4˚C
2. Wash with 80% Ethanol twice
3. Spin for 1 minute at 20,000g at 4˚C.
4. Remove residual EtOH, let pellet air dry until you confirm it visually.
5. Resuspend with 25 µL of TE
6. Measure with the Qubit
7. You can expect 10^6 colonies per electroporation. Perform more electroporations to get the desired coverage for your library. You should aim for a minimum of 50 fold coverage of your library.
8. Mix 10 ng of DNA with 50 µL of MegaX competent cells and leave on ice.
9. Electroporate using the following conditions: 2000V, 200 ohms, 25 uF in 0.1 cm Cuvette
10. After electroporation, add 1 mL of Recovery Medium to the cells in the cuvette and transfer the solution to a 15 mL culture tube. To recover residual cells in the cuvette, repeat this three more times to get a total of 4 mL
11. Shake at 225 rpm (37°C) for 1 hour
12. Spread about 660 µL onto one large LB plate (245 x 245 mm) with antibiotics. Total 6 plates will be used for one electroporation. To count total colony number, mix 1 or 2 µL with 200 µL of SOC media and spread on a 100mm plate with antibiotics.
13. Incubate plates for 20 hours at 37˚C to prevent colonies from overgrowing. The colonies should be visible, but small, and measure approximately 0.5mm in diameter. If too small, continue incubation, checking periodically.
14. After the colonies have grown up, compare the number of colonies in the full ligation and self-ligation transformations, and determine the background rate. We typically see a background rate < 0.5%.

**Step 8. Plasmid purification. (Day 6)**

1. After colonies have grown to an appropriate size, harvest colonies from one plate by scraping 3 times with LB containing Carb in the following order: 25 mL, 10 mL and finally 5 mL.
2. Collect pellets by centrifugation at 6000 rpm for 20 min.
3. Store all tubes in –80˚C or proceed to purify plasmids using the ZymoPure Plasmid Miniprep kit according to manufacturer's conditions.
4. Combine E. coli pellets from two large LB plates and use 3 maxi-prep columns for the purification of plasmid library. 9 Maxi-prep columns will be required for one electroporation.
5. Pool all the purified plasmid libraries and perform the QC with the Nanodrop to measure concentration, and store the plasmid library in aliquots at -20˚C.
6. Prepare NGS sequencing library by PCR reaction to determine the representation of sgRNAs.

#### Appendix:

**Table of adapters and primers to clone subpools (from the Jonathan S. Weissman Lab)**

Append the adapter pair to each end of the oligo sequence without reverse complementing

| | | | |
|---|---|---|---|
| Adapter_001 | ATTTTGCCCCTGGTTCTT | Adapter_001 | CCAGTTCATTTCTTAGGG |
| Adapter_002 | TCACAACTACACCAGAAG | Adapter_002 | GCAACACTTTGACGAAGA |
| Adapter_003 | TGAACTCCACCTCGTTAA | Adapter_003 | TTGCTTCCCCCATGCTTT |
| Adapter_004 | CTGTGTAATCTCCGACAC | Adapter_004 | GCCTTTGCATGTTGTGGA |
| Adapter_005 | GCGTGTTTGAATTCCACT | Adapter_005 | AAATTTCCTCGTCGGCTC |
| Adapter_006 | AGGATCTCTAGCCTCAAA | Adapter_006 | GATGAAGCATCGTAACTG |
| Adapter_007 | AACTGCGATCGCTAATGT | Adapter_007 | GTTCTCCAGTGCCTTATT |
| Adapter_008 | CTTCTACCGAACATACAG | Adapter_008 | TCGCGTTATGCTGTATGT |
| Adapter_009 | CGTAGCGTTTTGTACACG | Adapter_009 | CACGCTCAAAAAGCGTAC |
| Adapter_010 | AGTCCATCGCCGACATTA | Adapter_010 | AGCGAATTTGCGCTGACA |
| Adapter_011 | GGCATGCTGCAATAACCT | Adapter_011 | AAACCGGTGAGCTGGAAT |
| Adapter_012 | CCGGTAACTATTCTAGCC | Adapter_012 | GCAGCCCGAATACTTTCA |
| Adapter_013 | GTGATCCGCTCCACTTAA | Adapter_013 | ACAGTGTTGTGATCTTGC |
| Adapter_014 | ATACGTTAACCCGTATGG | Adapter_014 | TCTAGCCTACAATTCACG |
| Adapter_015 | CCCTAAGCATTCGCGAAA | Adapter_015 | ATTGTTAGCGCCACAATC |
| Adapter_016 | CATTTCTAGCCCTTCGAG | Adapter_016 | AACTCAACTTGGCAGGAA |
| Adapter_017 | GGATTGGACAAGCTAGTT | Adapter_017 | TAATACAAGGCCGCGTTG |
| Adapter_018 | GGGGGAAAAGGATCGATT | Adapter_018 | TCACGTGTAAACGATGCG |
| Adapter_019 | ACCCCAGTTGTGAATATC | Adapter_019 | TCGGATGACCCTAGAAAG |
| Adapter_020 | TATGAACCACTAAGGCGT | Adapter_020 | CGTAAAGTCTGCTGGTGA |
| Adapter_021 | GCTGGTGTTATGGTGGAA | Adapter_021 | AAGTCGTTTTTGGACCGC |
| Adapter_022 | GAGCACACACAAGAATGA | Adapter_022 | TAGCGACAACGTCACAAC |
| Adapter_023 | CACACTATGTCATCCGCA | Adapter_023 | CCTTTAGCAAGCAAAGCC |
| Adapter_024 | TAGTCAGAGAGTCGAGAG | Adapter_024 | GGTTTGGAGCCATTAGTT |
| Adapter_025 | GTAGACAAATGCTTGGAC | Adapter_025 | TCCCAGTCTAGTATGAGG |
| Adapter_026 | AAGGCCCAAGTCGCTTTT | Adapter_026 | CCGCCTTTTTCAAGTGAT |
| Adapter_027 | TTCCTTTCGTTCTCCTTC | Adapter_027 | GCGTTCATTGGTACCTAG |
| Adapter_028 | TCGATCCGGGAGTATACA | Adapter_028 | CCTACAAGAGTTCGACAC |
| Adapter_029 | ACATCCTGGTTACTTGGC | Adapter_029 | TGCTCTCGATCATAGCCT |

**Primers to amplify 1-29**

Lowercase overhangs correspond to the 5' and 3' most 4 bases contained within the sgRNA oligo sequence

| | | | |
|---|---|---|---|
| adapter_001_fwd | ATTTTGCCCCTGGTTCTTccac | adapter_001_rev | CCCTAAGAAATGAACTGGcagc |
| adapter_002_fwd | TCACAACTACACCAGAAGccac | adapter_002_rev | TCTTCGTCAAAGTGTTGCcagc |
| adapter_003_fwd | TGAACTCCACCTCGTTAAccac | adapter_003_rev | AAAGCATGGGGGAAGCAAcagc |
| adapter_004_fwd | CTGTGTAATCTCCGACACccac | adapter_004_rev | TCCACAACATGCAAAGGCcagc |
| adapter_005_fwd | GCGTGTTTGAATTCCACTccac | adapter_005_rev | GAGCCGACGAGGAAATTTcagc |
| adapter_006_fwd | AGGATCTCTAGCCTCAAAccac | adapter_006_rev | CAGTTACGATGCTTCATCcagc |
| adapter_007_fwd | AACTGCGATCGCTAATGTccac | adapter_007_rev | AATAAGGCACTGGAGAACcagc |

| | | | |
|---|---|---|---|
| adapter_008_fwd | CTTCTACCGAACATACAGccac | adapter_008_rev | ACATACAGCATAACGCGAcagc |
| adapter_009_fwd | CGTAGCGTTTTGTACACGccac | adapter_009_rev | GTACGCTTTTTGAGCGTGcagc |
| adapter_010_fwd | AGTCCATCGCCGACATTAccac | adapter_010_rev | TGTCAGCGCAAATTCGCTcagc |
| adapter_011_fwd | GGCATGCTGCAATAACCTccac | adapter_011_rev | ATTCCAGCTCACCGGTTTcagc |
| adapter_012_fwd | CCGGTAACTATTCTAGCCccac | adapter_012_rev | TGAAAGTATTCGGGCTGCcagc |
| adapter_013_fwd | GTGATCCGCTCCACTTAAccac | adapter_013_rev | GCAAGATCACAACACTGTcagc |
| adapter_014_fwd | ATACGTTAACCCGTATGGccac | adapter_014_rev | CGTGAATTGTAGGCTAGAcagc |
| adapter_015_fwd | CCCTAAGCATTCGCGAAAccac | adapter_015_rev | GATTGTGGCGCTAACAATcagc |
| adapter_016_fwd | CATTTCTAGCCCTTCGAGccac | adapter_016_rev | TTCCTGCCAAGTTGAGTTcagc |
| adapter_017_fwd | GGATTGGACAAGCTAGTTccac | adapter_017_rev | CAACGCGGCCTTGTATTAcagc |
| adapter_018_fwd | GGGGGAAAAGGATCGATTccac | adapter_018_rev | CGCATCGTTTACACGTGAcagc |
| adapter_019_fwd | ACCCCAGTTGTGAATATCccac | adapter_019_rev | CTTTCTAGGGTCATCCGAcagc |
| adapter_020_fwd | TATGAACCACTAAGGCGTccac | adapter_020_rev | TCACCAGCAGACTTTACGcagc |
| adapter_021_fwd | GCTGGTGTTATGGTGGAAccac | adapter_021_rev | GCGGTCCAAAAACGACTTcagc |
| adapter_022_fwd | GAGCACACACAAGAATGAccac | adapter_022_rev | GTTGTGACGTTGTCGCTAcagc |
| adapter_023_fwd | CACACTATGTCATCCGCAccac | adapter_023_rev | GGCTTTGCTTGCTAAAGGcagc |
| adapter_024_fwd | TAGTCAGAGAGTCGAGAGccac | adapter_024_rev | AACTAATGGCTCCAAACCcagc |
| adapter_025_fwd | GTAGACAAATGCTTGGACccac | adapter_025_rev | CCTCATACTAGACTGGGAcagc |
| adapter_026_fwd | AAGGCCCAAGTCGCTTTTccac | adapter_026_rev | ATCACTTGAAAAAGGCGGcagc |
| adapter_027_fwd | TTCCTTTCGTTCTCCTTCccac | adapter_027_rev | CTAGGTACCAATGAACGCcagc |
| adapter_028_fwd | TCGATCCGGGAGTATACAccac | adapter_028_rev | GTGTCGAACTCTTGTAGGcagc |
| adapter_029_fwd | ACATCCTGGTTACTTGGCccac | adapter_029_rev | AGGCTATGATCGAGAGCAcagc |

**PCR-free areas**

Ideally, a dedicated PCR and plasmid free area of the lab is available to setup amplification reactions to avoid contamination. This space is also useful when extracting genomic DNA from CRISPR screens and setting up NGS sample prep PCR. Below are a few guidelines for a PCR-free area.

1. If a separate room cannot be dedicated for a PCR and plasmid free area, set aside a lab bay as far from plasmid and post-PCR work.
2. A PCR workstation is also beneficial to keep out contamination.
3. Personnel should avoid entering the space if they have worked with plasmids or amplicons earlier in the day.
4. When entering the space, dedicated lab coats and new gloves should be donned to avoid bringing in contaminants.
5. Supplies and reagents should never come from parts of the lab where plasmid and PCR amplicons are present.
6. If you need to bring out a sample for analysis or quantification but need to work with that sample in the PCR-free area, bring out a small aliquot so potentially contaminated material is not brought back.
7. If a contamination even has occurred, dispose of any contaminated materials and supplies. Clean contaminated surfaces with a 1:10 dilution of bleach (0.5% final). Leave the bleach on the surface for at least 15 minutes and then clean the surface with 70% EtOH to remove the bleach reside.
